# Blood flow modeling reveals improved collateral artery performance during mammalian heart regeneration

**DOI:** 10.1101/2021.09.17.460699

**Authors:** Suhaas Anbazhakan, Pamela E. Rios Coronado, Ana Natalia L. Sy-Quia, Anson Seow, Aubrey M. Hands, Mingming Zhao, Melody L. Dong, Martin Pfaller, Brian C. Raftrey, Christopher K. Cook, Daniel Bernstein, Koen Nieman, Anca M. Pașca, Alison L. Marsden, Kristy Red-Horse

## Abstract

Collateral arteries are a vessel subtype that bridges two artery branches, forming a natural bypass that can deliver blood flow downstream of an occlusion. These bridges in the human heart are associated with better outcomes during coronary artery disease. We recently found that their rapid development in neonates supports heart regeneration, while the non-regenerative adult heart displays slow and minimal collateralization. Thus, inducing robust collateral artery networks could serve as viable treatment for cardiac ischemia, but reaching this goal requires more knowledge on their developmental mechanisms and functional capabilities. Here, we use whole-organ imaging and 3D computational fluid dynamics (CFD) modeling to identify the spatial architecture of and predict blood flow through collaterals in neonate and adult hearts. We found that neonate collaterals are more numerous, larger in diameter, and, even when similar in size/number, are predicted to more effectively re-perfuse an occluded coronary network when compared to adults. CFD analysis revealed that collaterals perform better in neonates because of decreased differential pressures along their coronary artery tree. Furthermore, testing of various collateral configurations indicated that larger, more proximal collaterals are more beneficial than many smaller ones, identifying a target architecture for therapeutic interventions. Morphometric analysis revealed how the coronary artery network expands during postnatal growth. Vessel diameters do not scale with cardiac muscle growth. Instead, the coronary tree expands solely by adding additional branches of a set length, a burst of which occurs during murine puberty. Finally, we compared mouse structural and functional data to human hearts. Surprisingly, fetal human hearts possessed a very large number of small, but mature, smooth muscle cell covered collaterals while angiogram data indicated adult patients with chronic coronary occlusions contained at least two. Comparing size ratios with modeled mouse data suggested low re-perfusion capabilities of the embryonic collaterals but higher functional benefits of those in diseased adults. Our unique interdisciplinary approach allowed us to quantify the functional significance of collateral arteries during heart regeneration and repair–a critical step towards realizing their therapeutic potential.

## Introduction

Cardiovascular disease, including coronary artery disease (CAD), is the leading cause of death worldwide^1^. Atherosclerosis causes coronary arteries to become partially or completely occluded, decreasing blood flow to the myocardium and jeopardizing cardiac muscle function and viability. Current treatments include percutaneous interventions and coronary artery bypass graft surgery, but these are highly invasive and a significant number are unsuccessful, especially in diffuse multi-vessel CAD, which calls for new treatments^2^. Humans and some other mammals can develop specialized blood vessels called collateral arteries that function as natural coronary bypasses.

Collateral arteries are defined as an artery segment directly bridging two artery branches without intervening capillaries, such that they directly provide blood flow distal to a coronary blockage. Although only a minority of adult humans have functionally significant collateral arteries, clinical observations indicate that they can successfully shunt blood around a stenosis to protect against myocardial ischemia and reduce the risk of cardiac death^3–6^. Thus, inducing collateral development could be a promising therapeutic approach for treating CAD^7^. However, a major roadblock to this goal is the severe lack of knowledge about collateral developmental mechanisms and their ability to restore blood flow.

While studies have characterized the presence or absence of native collateral arteries across different mammals^8^, mice are the most common model for investigating their function during cardiac injury, usually through surgically-induced myocardial infarctions (MI)^9–13^. Mice do not generally have pre-existing collateral arteries, but they can be observed in adults by 7 days post-MI when using vascular filling approaches, i.e. Microfil injection into the vasculature. This method detects 6-10 collaterals per adult heart at approximately 18 μm in diameter^10^. Genetic deletions in chemokine receptors that inhibit macrophages reduces collateral numbers^10^. Furthermore, mouse strains with decreased collateral development have genetic variants that lower *Rabep2* expression, which encodes a protein involved in VEGFR2 endosomal trafficking and signaling^14^. Thus, mice have been a useful model for understanding various aspects of collateral biology.

We recently used a different technique to identify collaterals—whole-mount immunofluorescence—coupled with lineage tracing and mouse genetics to identify the cellular and molecular mechanisms driving collateral development post-MI^9^. We found that, in the regenerating neonate heart, collaterals form post-MI when arterial endothelial cells migrate into the infarct zone in response to hypoxia-induced CXCL12 and coalesce into collateral arteries. This process was termed artery reassembly and did not occur in the non-regenerative adult heart, suggesting that the collaterals observed during vascular filling (described above) utilized a different mechanism. Exogenous CXCL12 application induced artery reassembly in adults to create collaterals up to 40 μm in diameter. Although these collaterals were positively correlated with heart regeneration and repair, and vascular filling methods established direct connections, a detailed description of how blood flows through these relatively small vascular connections is required to fully understand the functional capabilities and therapeutic potential of collateral arteries.

How structural parameters affect collateral hemodynamics in these injury models remains an unanswered question due to technical barriers of directly imaging blood flow. Clinical measurements of collateral flow rely on qualitative assessments from angiograms or indirect pressure measurements^15–17^. More accurate measurements of collateral flow in humans are not only invasive, but somewhat unreliable since conclusive relationships cannot be made without knowing the number and size of all collaterals, many of which cannot currently be imaged in the human heart via angiogram. Visualizing blood flow is even more difficult in experimental animals due to their small size. Conclusions regarding collateral flow are usually reached from *ex-vivo* data, but not without significant limitations. Methods include: 1. Filling coronary vessels through the aorta (Microfil casting, µCT and fluorescent conjugates)^18–23^, which creates a non-cell specific volumetric map of coronary vessels, and 2. Whole mount immunostaining fixed hearts from postnatal transgenic mice^9^. None of these approaches provide a precise picture of how flowing blood will distribute through collaterals.

Because the physical laws governing fluid motion are known, computational fluid dynamics (CFD) modeling tools can directly and precisely estimate blood flow. CFD has contributed to patient-specific surgical and treatment planning in numerous human cardiovascular diseases^24–28^. CFD modeling has also been applied to the cardiovascular systems of various animals to estimate hemodynamic forces that would otherwise be difficult to directly measure^29–32^. In mice, computational studies have modeled blood flow in the retinal vasculature, thoracic aorta, and even feto-placental vessels for the purpose of defining how hemodynamic forces influence arterial remodeling events at the cellular and molecular levels^33–35^. An additional major advantage of CFD modeling is the ability to systematically alter certain parameters while keeping others constant, leading to rapid conclusions on the reparative capabilities of different vascular architectures^36^.

To obtain correct estimates from CFD modeling, it is critical to have high-resolution images of a vascular network with intact volumetric dimensions. To date, the vascular labeling methods utilized have not had the resolution required to generate detailed anatomic models suitable for CFD modeling. However, recent innovations in tissue clearing and whole-organ microscopy now provide the possibility of generating sufficiently high resolution images suitable for CFD model building^37–40^. Thus, CFD is perfectly poised to push forward our understanding of collateral function.

In this study, we sought to interrogate how different collateral configurations affect blood flow post-injury. We optimized whole-organ immunostaining and clearing to label and image the entire intact artery tree, allowing quantification of hemodynamic forces via CFD using high-fidelity 3D models constructed in the neonate and adult. We computationally generated virtual occlusions and various collateral configurations, keeping other model parameters fixed, to measure levels of flow restoration. The results showed that naturally forming collaterals in adult mouse hearts perform poorly and restore little flow. The virtual equivalent of CXCL12-induced collaterals performs better but remain sub-optimal. In contrast, naturally forming collaterals in neonate hearts are highly restorative because the structural parameters of the early coronary tree and cardiac output at this stage results in lower overall pressure loss along the coronary tree. We additionally investigated collateral arteries in human hearts by generating whole-organ images of fetal hearts and analyzing angiograms from patients with chronic coronary occlusions. Surprisingly, we found both fetal hearts contained greater than 40 mature, smooth muscle covered collateral arteries, while only an average of two collaterals with measurable flow in patient angiograms. Comparing diameters to CFD mouse models where flow restoration was quantified suggested that human fetal collaterals may be too small to be functionally significant and that those in patients may lie between the capabilities of neonate and adult mouse collaterals. In total, the combination of whole-organ artery labeling with 3D CFD modeling provides a powerful tool to accurately analyze hemodynamic forces in collateral arteries to broaden our understanding of their functional significance and therapeutic potential.

## Results

### Imaging the entire coronary artery tree in neonate and adult mouse hearts

To utilize 3D CFD for modeling coronary blood flow at high resolution and with controlled parameter perturbations, we required a method to image the entire intact artery tree in three dimensions. A whole-organ immunostaining and clearing method based on iDISCO was optimized for cardiac tissue using postnatal day 6 (P6) mice^37, 39^, which allowed us to image smooth muscle actin (α-SMA)-positive arterial smooth muscle cells throughout the heart using Light sheet microscopy (Fig. 1a). The signal-to-noise ratio of α-SMA staining was high, allowing sharp contrast of arteries throughout the entire myocardium (Fig. 1b). Rendering the arterial immunolabeling in 3D using Imaris software revealed vast improvements over our previous method^9^. Specifically, the 3D architecture was retained (Fig. 1c), and we could fully observe the septal artery (SpA) in addition to the left (LCA) and right (RCA) coronary arteries (Fig 1c_ii_). Another improvement was the ability to fully immunolabel and image intact adult hearts (Fig. 1d). Immunostaining of α-SMA labeled arterial vessels throughout the entire adult heart (Fig. 1e), even deep within the septum (Fig. 1f). Co-staining with arterial endothelial marker, Connexin40, confirmed extensive overlap in both neonate and adults (Fig. 1b_iii-v_ and data not shown). Qualitatively, the density of arteries is much greater in neonates compared to adult (Compare Fig.1a and 1d), and more branches were detected when compared to µCT methods, both in neonatal and adult stages^41^. These data demonstrate that iDISCO and Light sheet microscopy together, are capable of effectively labeling and imaging smooth muscle covered arteries throughout neonatal and adult hearts.

**Figure 1:**
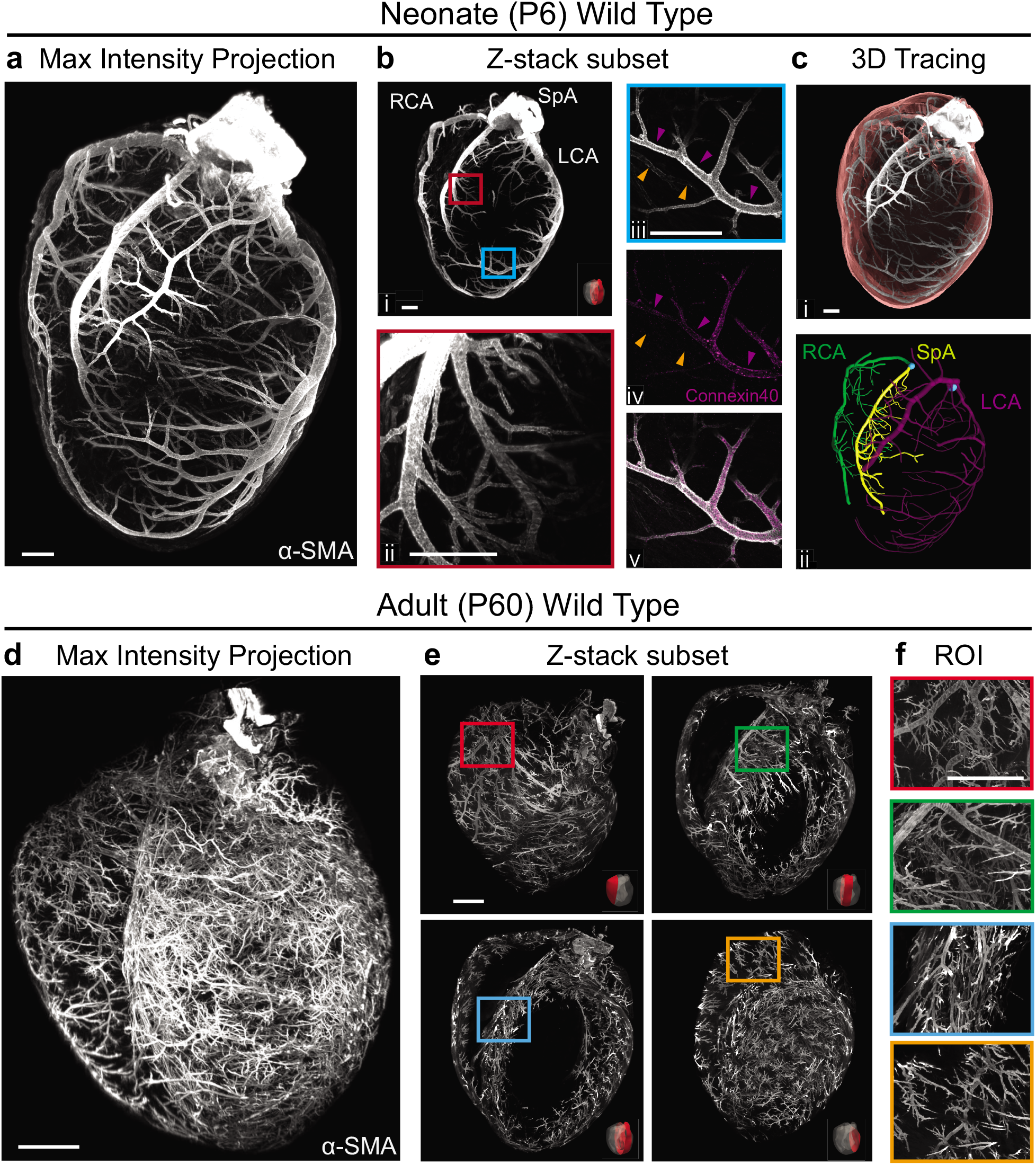
Whole-organ imaging of coronary arteries at cellular resolution. Neonate and adult hearts (atria removed) subjected to tissue clearing and immunolabeling with alpha-smooth muscle actin (α-SMA) and imaged using a Light sheet microscope. (**a**) Maximum intensity projection of entire neonate heart. (**b**) Z-Stack subset projections of Light sheet images (**b_i-ii_**) or those captured using a confocal microscope (**b_iii-v_**). High magnification of septum shows complexity of the septal artery (**b_iii_**) and of LCA shows colocalization of α-SMA+ branches with artery marker Connexin40 (purple arrowheads)(**b_iii-v_**). An α-SMA^low^Connexin40-vein (orange arrowheads) is also present in **b_iii-v_**. (**c**) 3D rendering of myocardial volume (red)(**c_i_**) and main coronary artery branches: Right (RCA), Septal (SpA) and Left (LCA)(**c_ii_**). (**d**) Maximum intensity projection of entire adult heart. (**e** and **f**) Z-Stack subsets of indicated heart regions (**e**) and region-of-interest (ROI) images (**f**) reveal the high resolution and specificity of immunolabeling with this technique. Scale bars: **a-c**, 300 μm; **d-f**, 500 μm.

To characterize collateral arteries using this novel method, we imaged neonatal and adult mice subjected to myocardial infarction (MI)(uninjured mouse hearts do not generally contain collaterals)^9, 10^. Collateral arteries form faster in neonates than in adults^9^. Thus, injured neonatal hearts were harvested 4-days post MI while adult hearts were collected 28-days post-MI, followed by arterial immunolabeling and clearing. A collateral tracing pipeline was developed first using images of neonatal hearts (Fig. 2a). ImageJ’s Simple Neurite Tracer plugin^42, 43^ was used to trace, in a semi-automated way, every α-SMA+ vessel that originated downstream of the LCA occlusion (suture) and connected to either the RCA, SpA, or the LCA upstream of the occlusion. Traced paths were isolated and masked so that 3D rendering in Imaris created a map of every collateral artery found post-MI (Fig. 2b). The resolution of our method allowed us to annotate the precise collateral segments that bridged two artery branches (Fig. 2b_ii_). A collateral bridge was defined as the segment of continuous smooth muscle covered vessel that existed between two branch tips with opposing branch angles (Fig. 2c). Tracing did not detect collateral connections in non-injured neonate hearts (Fig. S1). Thus, this method reliably identifies collateral arteries in whole heart images.

**Figure 2:**
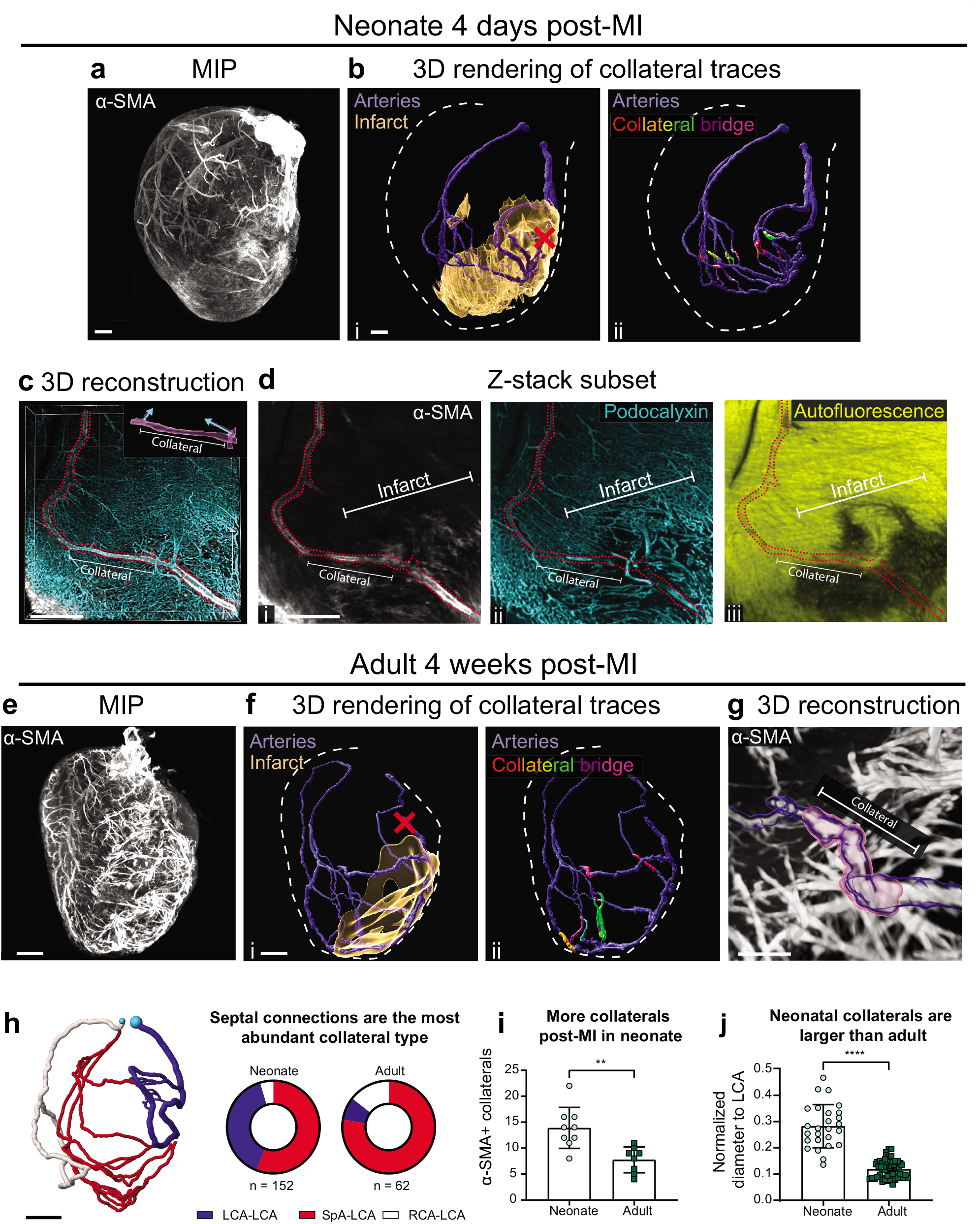
Increased collateral arteries in neonate versus adult hearts post injury. (**a-d**) Whole organ imaging of P6 neonatal heart labeled with α-SMA post myocardial infarction (MI). (**a**) Maximum intensity projection (MIP) of entire heart. (**b**) Collateral connections traced from downstream of suture (red X) were 3D rendered and overlaid with infarct volume (**b_i_**) and collateral bridges (**b_ii_**). (**c**) 3D reconstruction of 100 μm Z-stack containing a representative collateral bridge within a traced vessel (red dotted line). (**d**) MIP of a 35 μm Z-stack within **c** highlighting an α-SMA+ collateral (**d_i_**) and its relation to Podocalyxin labeling all vessels (**d_ii_**) and Autofluorescence labeling surviving myocardium (**d_iii_**). (**e-g**) Adult (16-week-old) injured hearts labeled with α-SMA. (**e**) MIP of entire heart. (**f**) 3D rendering of collateral connections overlaid with infarct volume (**f_i_**) or collateral bridges (**f_ii_**). (**g**) 3D reconstruction of representative collateral bridge (pink). (**h**) Classification and distribution of collateral connections. (**i**) Collateral numbers in neonate (n=9 hearts) and adult (n=8 hearts) post-MI. (**j**) Collateral diameters in neonate (n=26 hearts) and adult (n=55 hearts) post-MI. Scale bars: **a-b** and **e-f**, 300 μm; **c-d** and **g-h**, 150 μm. Right (RCA), left (LCA), and septal (SpA) coronary arteries. Error bars are st dev: **, p≤.01; ****, p≤.0001.

To ascertain where collateral bridges were localized with respect to injured myocardium, we labeled all coronary vessels in the neonate with Podocalyxin and used the autofluorescence signal to observe surviving cardiac muscle. Areas lacking autofluorescence, which were not present in uninjured hearts, delineated injured myocardium, which was confirmed by accompanying disrupted vasculature (Fig. 2d). Injured regions were outlined and overlaid onto collateral models (Fig. 2b_i_). Collateral bridges were usually located at the edge of the infarcted area, connecting regions of muscle and vascular death to unaffected sites in the heart (Fig. 2d). These same methods were then used to identify collaterals and injured myocardium in adult hearts. Similar patterns were observed (Fig. 2e-g).

We next quantified collateral connection type, numbers, and relative sizes in our images. Collateral connections were categorized based on which artery they connected (Fig. 2h). The majority of connections in both neonate and adult hearts were SpA-LCA. Neonate hearts formed more LCA-LCA and fewer RCA-LCA connections than adults (Fig. 2h). Neonate hearts also formed approximately 40% more collaterals than adults (Fig. 2i), and their diameters were larger relative to the proximal LCA (Fig. 2j). These data highlight the importance of advanced imaging methods for observing accurate vascular remodeling patterns, i.e., those involving the septal artery, and underscore the significant differences between young and old hearts.

### Modeling coronary blood flow

We next sought to understand how these collaterals might restore blood flow in the presence of a vascular occlusion. An *in silico* approach was employed that would allow us to computationally estimate blood flow while at the same time manipulating different parameters in isolation, such as collateral number, size, and location. First, an anatomically representative model of the native adult coronary tree was created using the open-source software, SimVascular (www.simvascular.org)^44^, from a Light sheet image of a non-injured adult heart labeled with α-SMA (Fig. 3a). The Light sheet images (Fig. 3a_i_) were used as a guide for drawing path lines through every artery in the heart up to tertiary branches (Fig. 3a_ii_, Methods). Arteries were then segmented by drawing a circle that encompassed the entire width at even intervals along the vessel (Fig. 3a_iii_). SimVascular was used to convert the segmentations into a 3D model (Fig. 3a_iv_). We next measured the amount of tissue shrinkage that occurs during iDISCO by calculating heart volumes pre- and post-clearing (Fig. 3b_i_). Shrinkage was on average 37% (Fig. 3b_ii_), and, thus, the model was computationally uniformly scaled up by 1.58-fold (Fig. 3b_iii_). The result was a model reflecting the realistic anatomic 3D architecture of an adult mouse coronary artery tree.

**Figure 3:**
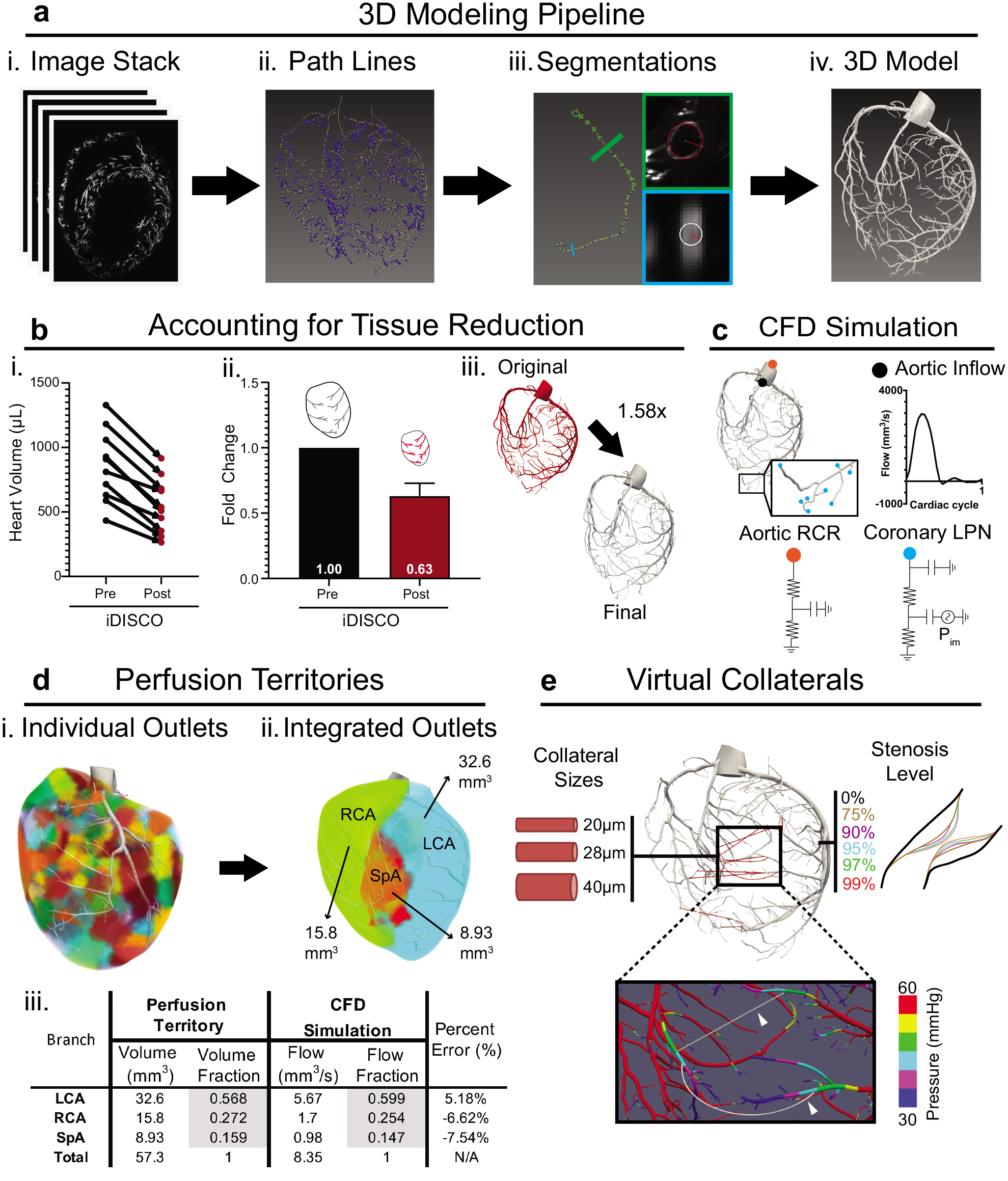
Building a physiologically representative 3D model of mouse coronary arteries. (**a**) Pipeline for generating 3D models from Light sheet images. (**b**) Scaling model to account for tissue volume reduction during iDISCO procedure. Measuring heart volumes pre- and post-processing (**b_i_**) yielded an average reduction value (**b_ii_**) used to generate a scaling factor for models (**b_iii_**). (**c**) Schematic of coronary simulation with a prescribed flow waveform at the inlet, RCR boundary condition at the aortic outlet, and coronary LPN at each coronary outlet. (**d**) Determining perfusion territories required first utilizing the Voronoi algorithm to outline perfusion subvolumes for each individual outlet (**d_i_**). Then, subvolumes were grouped by right (RCA), left (LCA), and septal (SpA) coronary branches (**d_ii_**). Outlet boundary conditions were tuned by matching simulated flow splits to perfusion territories (**d_iii_**). (**e**) Schematic depicting variations on collateral and stenosis parameters used in this study. Collaterals were placed to connect approximately equal pressure zones (white arrows). RCR, 3-element Windkessel model; LPN, lumped parameter network; *P_im_*, intramyocardial pressure.

This model was then used to computationally estimate physiologically realistic blood flow parameters throughout the arterial network. The simulations first required setting boundary conditions. At the aortic inlet, a flow waveform was set based on experimentally measured blood velocities from the literature for the neonate^45^ and adult^46^. Two outlet boundary conditions were set: 1. An RCR Windkessel model representing the systemic circulation at the aortic outlet^47^, and 2. A lumped parameter network (LPN) representing the coronary vessels downstream of the 3D model^48, 49^ (Fig. 3c). The lumped parameters included values accounting for vessel resistance at downstream arteries, capillaries, and veins and intramyocardial pressure due to contraction of the ventricle (Fig. 3c). Simulations were run on initial estimated parameters (see Methods) and were subsequently tuned to match expected flow splits between coronary branches to ensure our CFD simulation was distributing the flow proportionally. Flow splits were calculated based on perfusion territories for each of the 3 main branches of the coronary arteries. Each region of the myocardium was connected to its closest arterial end branch, and all the subregions were identified as belonging to branches of either the LCA, RCA, or SpA (Fig. 3d_i-ii_). The method estimated the LCA, RCA, and SpA to perfuse 60%, 25%, and 15% of the myocardium, respectively (Fig. 3d_ii_). Using this information to tune outlet boundary conditions (Fig. S2, Methods) resulted in close agreement between estimated perfusion territory and simulated flow splits (Fig. 3d_iii_). In total, the adjustments to the model and boundary conditions provided a model with close concordance to native physiology.

Our next goal was to investigate collateral blood flow, and one benefit of a computational approach is that parameters, such as collateral number/size and stenosis severity, can be virtually modified and systematically tested (Fig. 3e). We placed virtual collaterals within the native coronary tree model described above, using post-injury imaging data to guide general placement (see Fig. 2). Computationally derived pressure values were then used to precisely adjust placement at each branch so that collaterals joined two regions of equal pressure. This minimized flow across collaterals without stenosis, which is important to establish a consistent baseline so that different configurations could be properly compared (Fig. 3e, Fig. S3). These guidelines were used to produce 5 different collateral configurations in the adult heart (Fig. S3). We compared pressure difference, flow, and shear stress in all collaterals from each configuration to Poiseuille’s law, which analytically describes flow through a circular cylinder, to ensure the results of our simulations were reasonable (Fig. S4, Methods). These data verified virtually-placed collaterals for use in computational flow modeling.

Because we wanted to make comparisons between adult and neonate hearts, we performed the same workflow with an uninjured P6 heart. Perfusion territories were similar, but a lower aortic inflow was prescribed for neonates to match lower mean pressures following published values^45, 50^. Four collateral configurations were produced for neonates (Fig. S3). Then, adult and neonate models were used to investigate re-perfusion upon virtual stenosis.

### Investigating flow recovery by collateral arteries

We next sought to understand the level of flow recovery by virtual collaterals in the presence of coronary occlusions. One way to quantify re-perfusion is to sum the flows from all outlets downstream of the virtual stenosis and compare this to a normalized baseline flow with no stenosis (set at 100%). As expected, with no collaterals in adults total flow downstream of stenosis decreases when percent occlusion increases, especially above 90% (Fig. 4a, top row in chart). When comparing all the configurations tested at all stenosis levels, the configuration with 9 collaterals at 40 μm (9col, 40μm) provides the most flow recovery benefit, especially at 99% stenosis where it restores almost 25% of the non-stenotic flow compared to just 1% without collaterals (Fig. 4a, right-most column in chart). However, this extent of collateralization does not occur naturally with coronary artery ligation in adult mice (see Fig. 2i and refs 9,10). We noted that configurations similar to those observed experimentally, i.e. 6-12col, 20μm, recovered very little flow as measured by this method (Fig. 4a). These data demonstrate that collateral arteries as they naturally form after adult coronary occlusion are not expected to appreciably recover blood flow, but that increasing diameters, which is a major factor in reducing overall resistance, could enhance their function.

**Figure 4:**
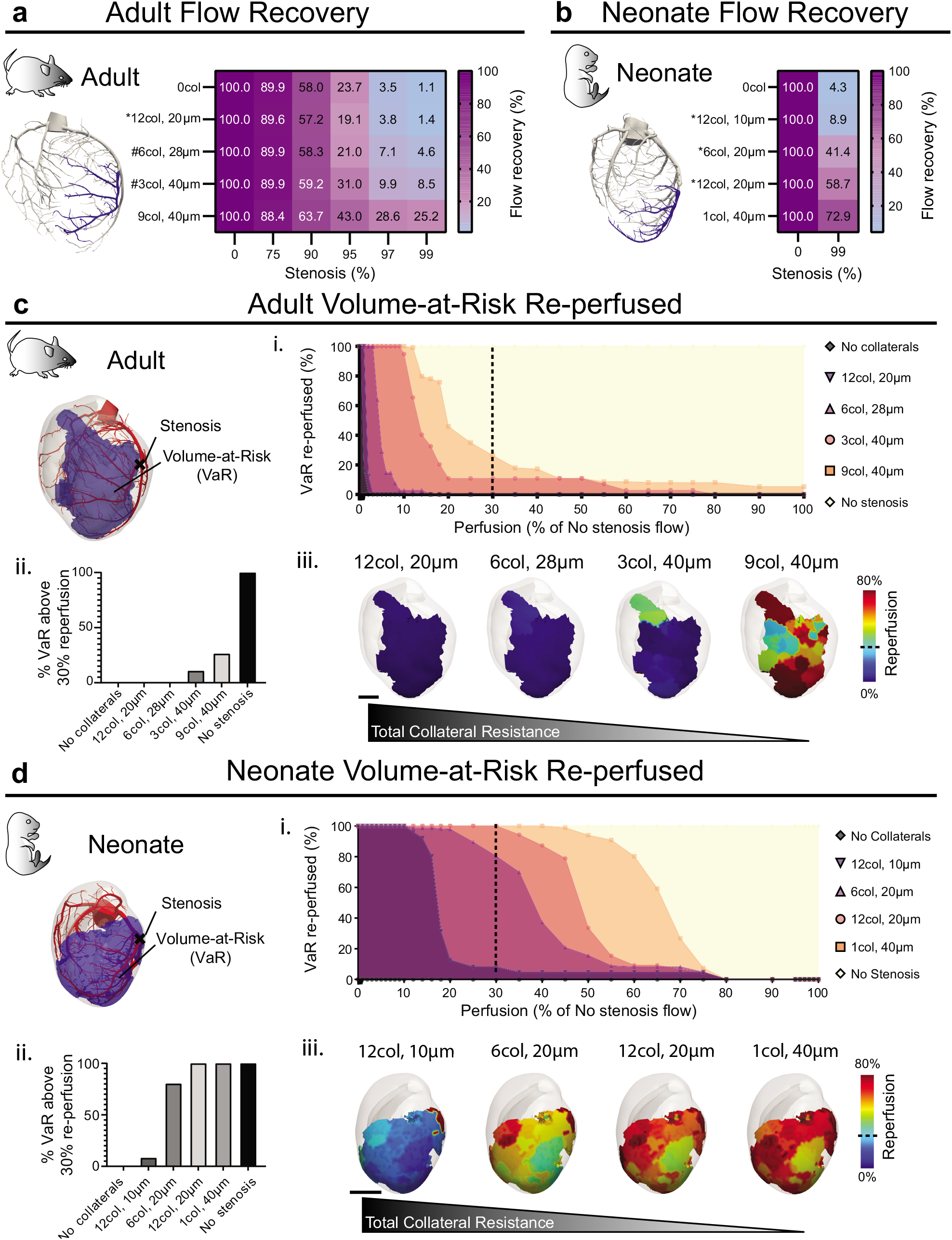
Collateral arteries are predicted to perform better in neonate hearts. (**a** and **b**) Measuring re-perfusion capacity by calculating percent of total non-stenotic flow in vessels downstream of the virtual occlusion (blue vessels). Asterisks denote configurations observed experimentally; hashtags denote sizes observed following CXCL12 injection^9^. Functionally significant re-perfusion is only seen in neonates under physiological conditions. (**c** and **d**) Percent re-perfusion of myocardial volume-at-risk (VaR)(blue region) in adult (**c**) and neonatal (**d**) models. Cumulative histogram (**c_i,_ d_i_**), bar graph of percent VaRs above 30% (**c_ii,_ d_ii_**), and visual maps of re-perfusion within the VaR (**c_iii,_ d_iii_**). Dotted line marks the ischemia threshold of 30% non-stenotic flow. Scale bar: 1000 μm.

In contrast to the poor function of adult collaterals, those of the same size and number in neonates performed well. The configuration that naturally forms in neonates (i.e. 12col, 20μm, see Fig. 2i and ref 9) is estimated to recover up to 60% of total flow downstream of a 99% stenosis (Fig. 4b). Remarkably, the largest diameter tested (40 μm) only required one vessel to provide massive recovery in neonates (Fig. 4b, last row in chart). As mentioned above, to compare adult and neonate flow recoveries, it is important to confirm that collaterals generally connect equal pressure zones (+/-10 mmHg) so that all configurations start with a similar collateral flow. This was further evident by the observation that adding collaterals did not change total downstream flow without stenosis and primarily increased flow only with increasing stenosis severity (Fig. S3). We concluded that collaterals in neonate hearts perform better than in adults.

The above analysis calculated overall recovery of pre-stenosis levels, but clinical data indicate that myocardial tissue could be supported at approximately 30% of baseline flow^51, 52^. Thus, we next sought to gain a more nuanced understanding of recovery by considering individual outlet perfusion territories downstream of the stenosis, so that we could observe if certain regions were receiving sustainable re-perfusion (i.e. >30% re-perfusion). First, we grouped all perfusion territories downstream of the stenosis to obtain the full volume-at-risk (Fig. 4c) and then plotted the percentage of that volume that is re-perfused above a certain threshold (Fig. 4c_i_). This revealed that while there were still no sustainably re-perfused regions in the 12col, 20μm and 6col, 28μm configurations, the 3col, 40μm and 9col, 40μm configurations were able to sustain 10 and 25% of the volume-at-risk, respectively (Fig. 4c_i-iii_). However, in the neonate, the 6col, 20μm re-perfused 80% of the volume-at-risk over the 30% threshold while the 12col, 20μm and 1col, 40μm configurations re-perfused the entire volume-at-risk (Fig. 4d_i-iii_). These data emphasized that collateral configurations of the same size, and thus same resistance, function better in the neonatal heart.

We next explored whether a more favorable placement of collaterals could improve the poor performance seen in adults. We started with a 3col, 40μm configuration (Fig. 5a, blue) and moved each collateral to a more proximal location in the coronary tree (Fig. 5a, red). This manipulation almost doubled total flow recovery (Fig. 5b) and approximately tripled the volume of myocardium re-perfused above the 30% threshold (Fig. 5c-e). Thus, variation in location can improve collateral function, likely because the pressure difference with more proximal attachments is much greater, resulting in increased flows.

**Figure 5:**
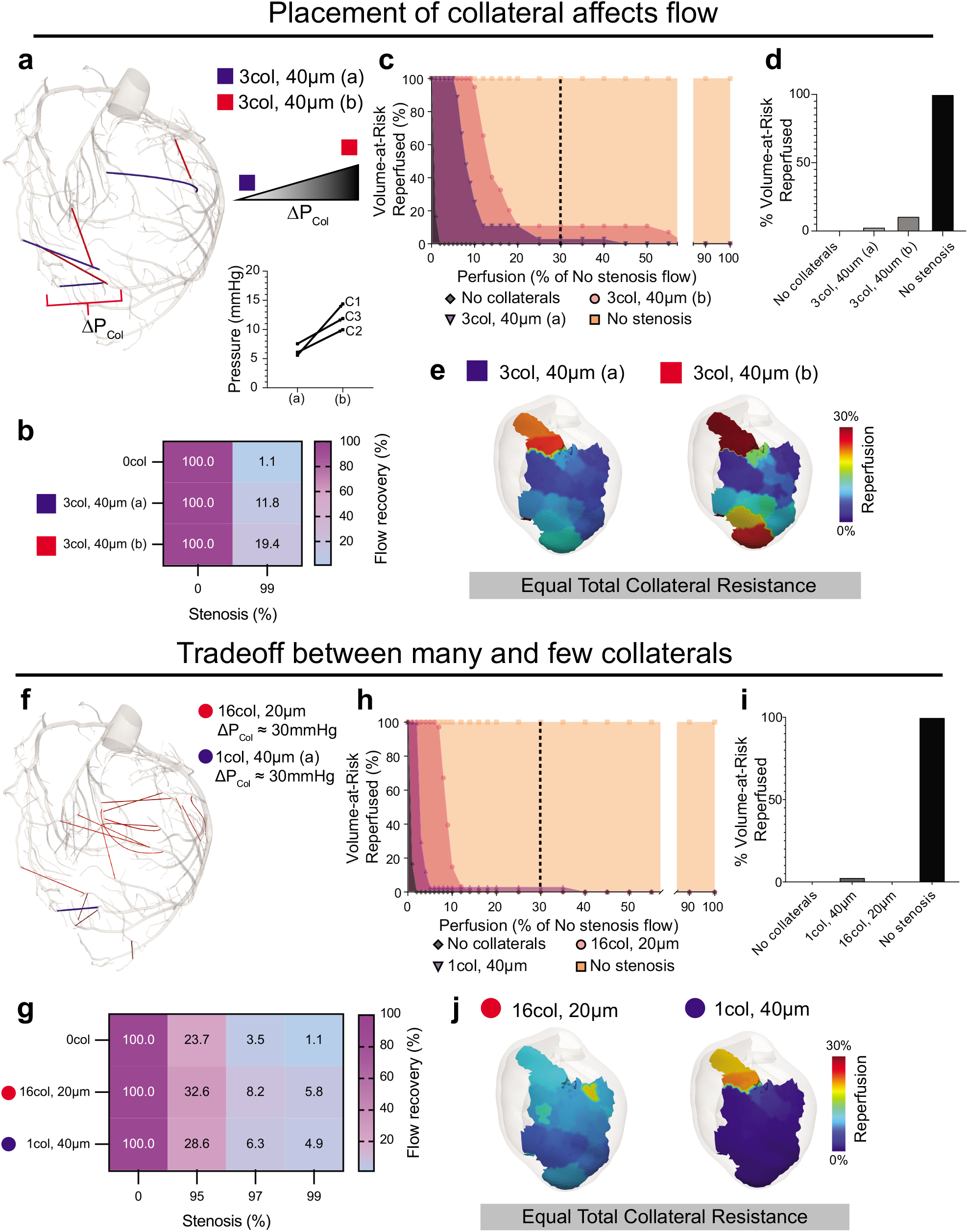
Evaluating collateral placement and the tradeoff between collateral number and size. (**a-e**) Investigating how collateral placement affects re-perfusion using two collateral configurations. (**a**) 3 collaterals with either high (red square) or low Δ*P_Col_* (blue square). (**a**) Graph showing the Δ*P_Col_* changes caused by altering placement for each collateral. (**f-j**) Investigating re-perfusion tradeoff between many, small and few, large collaterals using two configurations: 16 collaterals at 20μm (red circle) and 1 collateral at 40μm (blue circle). (**b, g**) The total non-stenotic flow in vessels downstream of the virtual occlusion. Cumulative histogram (**c, h**), bar graph of percent volume-at-risk (VaR) above 30% (**d, i**), and visual maps of re-perfusion within the VaR (**e, j**). Dotted line marks the ischemia threshold of 30% non-stenotic flow. Δ*P_Col_*, pressure difference across collaterals.

The above data suggested that fewer, larger collaterals are better than many, smaller ones (see Fig. 4). However, in those experiments, the total collateral resistance varied between the configurations. We tested this hypothesis by varying the number and size of the collaterals while keeping the total resistance equal. Simulations were performed on 2 configurations—16col, 20μm and 1col, 40μm (Fig. 5f). While total flow recovery was approximately equivalent (Fig. 5g), the 1col, 40μm configuration was uniquely able to re-perfuse 5% of the volume-at-risk above 30% (Fig. 5h-j). This analysis shows that fewer, larger collaterals could be more beneficial because they at least protect a portion of the myocardium while many, smaller collaterals distribute the re-perfusion so that none reach protective levels.

### Adult vs. neonate coronary artery morphology

Given that collaterals of the same size and total resistance were predicted to proportionally recover more flow in the neonate, we sought to understand why and first investigated arterial pressures at collateral formation sites at both ages. To facilitate comparisons between the two time points, Strahler ordering was used to classify branch segments into orders based on hierarchal position in the coronary tree and vessel diameter^53, 54^. Order 13 represented the aorta, order 12 represented the most proximal coronary artery segments, and subsequent orders represented downstream vessels until 8, which were the most distal branches modeled (Fig. 6a). This was used to compare hemodynamic and anatomical quantities at similar points in the coronary tree in both the adult and neonate. While absolute aortic and proximal coronary (order 12) pressures were vastly increased in the adult, the pressures at the most distal coronary tips (order 8) were approximately equal (Fig. 6b and c). Quantification revealed that the pressure drop along the coronary tree was ∼20 and ∼50 mmHg in neonate and adults, respectively (Fig. 6c). This is also true when considering just the segments downstream of the stenosis, making ΔP_Ad_ > ΔP_Neo_ (Fig. 6b and c). Thus, the collateral pressure difference (ΔP_Col_) required to restore pre-stenotic flow downstream of the occlusion is higher in the adult. Specifically, the ΔP_Col_ needs to be about ∼2-fold more in the adult to restore the same flow. Given that we see similar distal pressures at both stages, this explains why, even though collaterals in both recover the same absolute flow, it is much lower than the baseline, non-stenotic flow in the adult.

**Figure 6:**
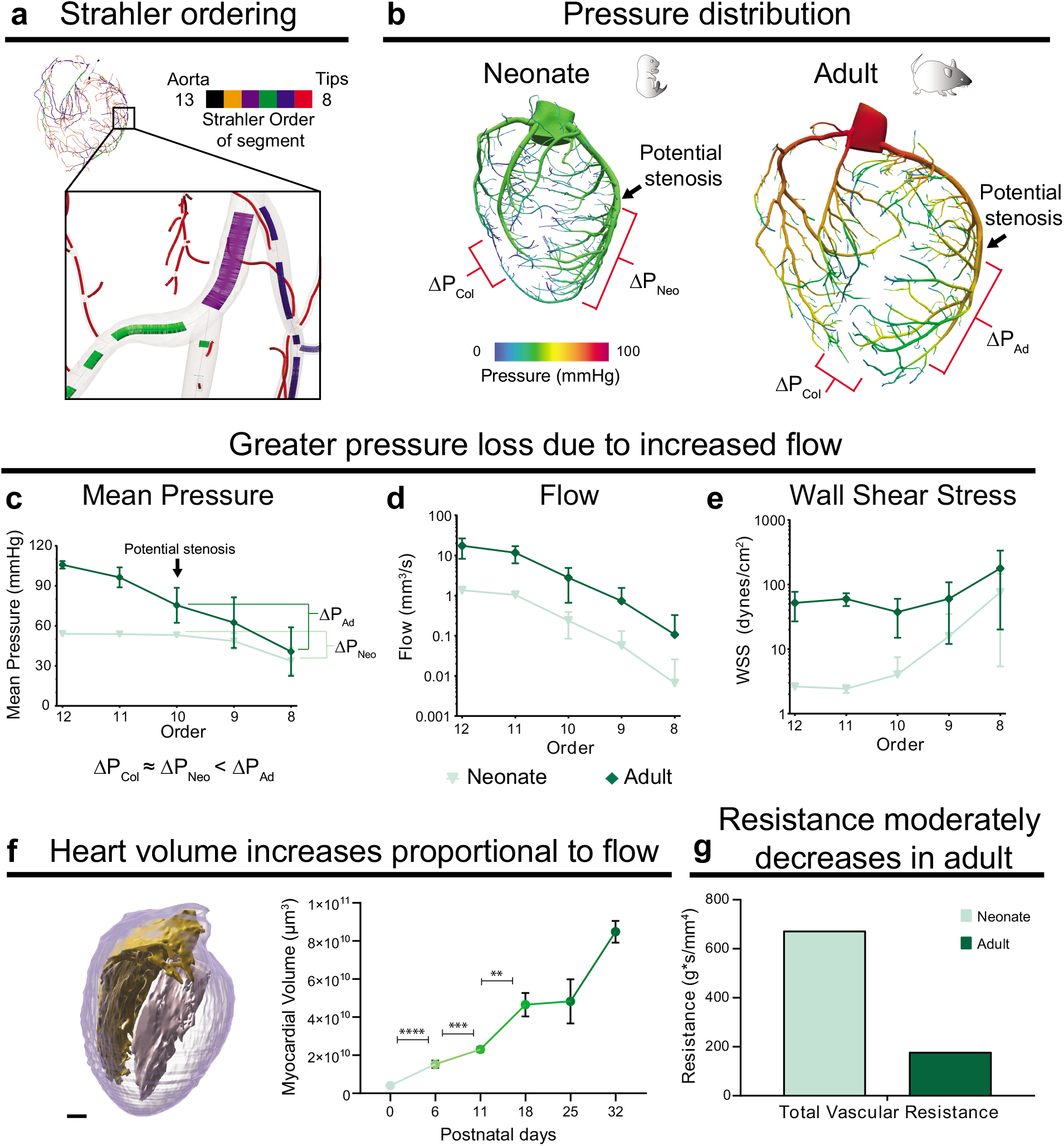
Investigating hemodynamic differences between neonate and adult. (**a**) Strahler ordering categorizes segments of the arterial tree from order 13 (aorta) to order 8 (distal tips). (**b**) Pressure distribution in the neonate and adult coronary models. (**c-e**) Quantification of pressure (**c**), flow (**d**), and wall shear stress (**e**) vs. Strahler order (n=1 P6, n=1 P60 heart model). (**f**) Heart volume segmentation (left) and quantification (n=3 P0, n=7 P6, n=3 P11, n=2 P18, n=2 P25, n=2 P32 hearts). (**g**) Total 3D resistance of the coronary tree in neonate and the adult models revealed a 3-fold decrease in adults. Δ*P_Col_*, pressure difference across collaterals; pressure difference downstream of a potential stenosis in the neonate, Δ*P_Neo_*, and the adult, Δ*P_Ad_*. Error bars are st dev: **, p≤.01; ***, p≤.001; ****, p≤.0001.

Our next experiments were aimed at understanding why ΔP_Ad_ was greater than ΔP_Neo._ Two factors critical for determining ΔP are flow rate and total resistance of the coronary tree. First, we compare the flow rate at each Strahler order between the neonate and the adult coronary models. Literature values indicated that aortic flow in adults is approximately ten times more than neonate, which was used as the inflow boundary condition for the computational model (see Fig. 3c)^45^. Simulations revealed that flow was also 10-fold greater for every vessel order modeled in the coronary tree (Fig. 6d). Shear stress was lower in neonates compared to adults, particularly in higher order vessels (Fig. 6e). We confirmed this trend held true when increasing the mesh size from 1.8 to 10 million elements; there was less than 10% difference in average shear stress with increased mesh resolution. Flow values were in line with increases in myocardial volume over time, i.e. volumes at P32 were more than 10-fold of P0 (Fig. 6f). The ability to rapidly determine volumes allowed us to analyze additional timepoints, which revealed a linear increase in myocardial volume during the first two weeks of life, a plateau between P18-25, and a burst of growth from P25-32. Second, we used the simulated flows and pressures to calculate the total resistance of the 3D coronary model. Neonate total vascular resistance was 3-fold that of adults (Fig. 6g). Since flow was increased by 10-fold, the 3-fold decrease in total resistance is not enough to offset flow increases. Thus, while the resistance of the coronary vasculature decreases in the adult, it’s not able to lower the resistance enough to balance the much greater flow, which manifests in a greater ΔP in adults.

We next investigated what features contribute to the nonproportional decrease in resistance with respect to the flow increase from neonate to adult. Two factors critical for determining resistance are vessel diameter and number of branches. Increases in these parameters both work to lower total resistance, diameter being the most impactful. Surprisingly, we found that the diameters were the same across all Strahler orders in each model (Fig. 7a). We validated this by comparing diameters of the most proximal segments of the RCA, SpA, LCA and aorta in multiple replicates of neonatal and adult hearts (Fig. 7b). The coronary stem diameter remains virtually the same while aortic diameter increased with age (Fig. 7b), a result we validated using an orthogonal method (Fig. 7c). Thus, coronary diameters do not grow proportionally to heart volume, which suggests that diameter expansion does not function to relieve vascular resistance in the face of increased flow demand in adults.

**Figure 7:**
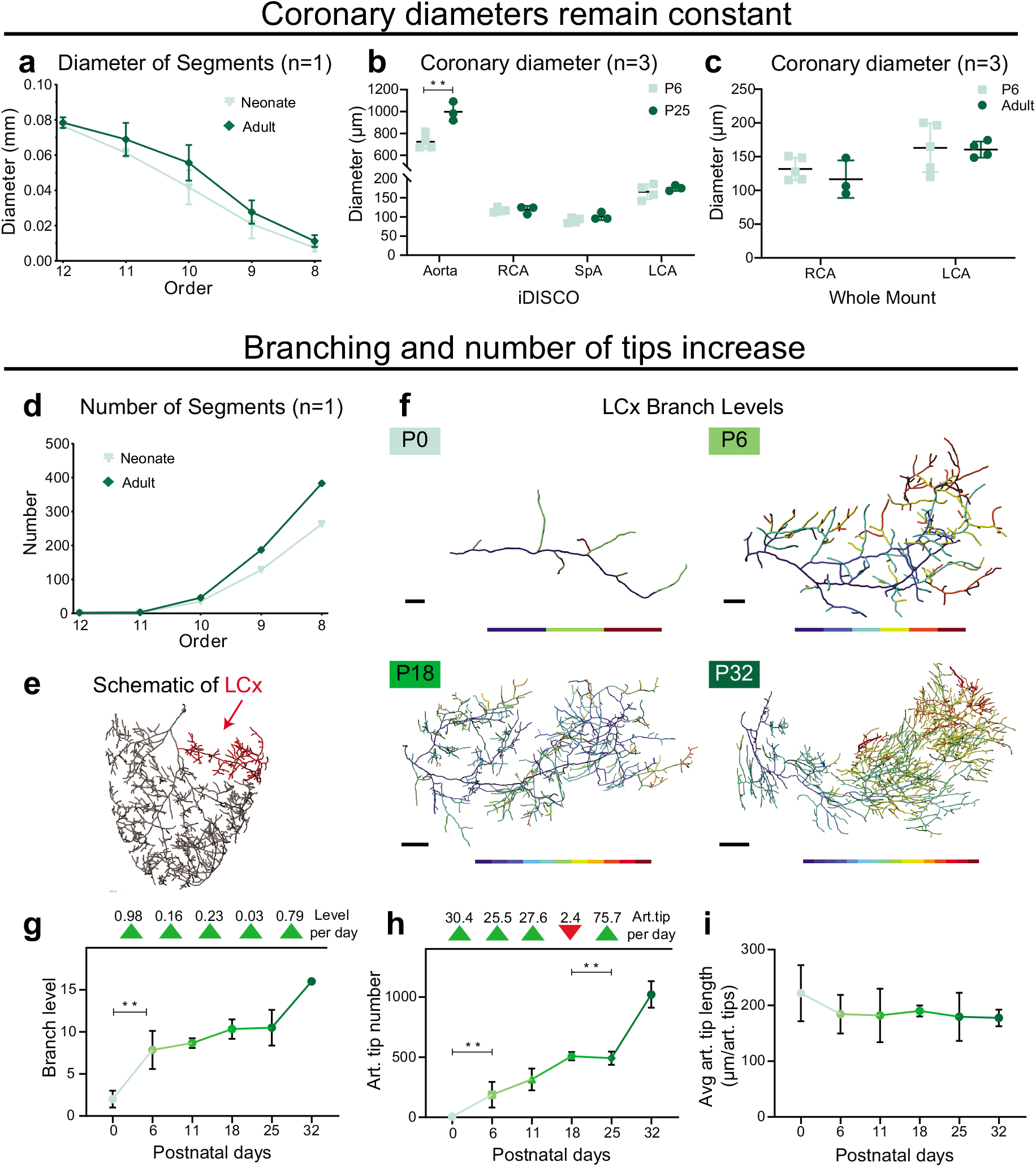
Main branch coronary artery diameters remain constant while branching increases throughout postnatal development. (**a**) Quantification of diameter vs. Strahler order. (**b** and **c**) Main branch coronary diameter measurements from additional hearts processed through iDISCO (**b**) or conventional whole mount immunostaining without clearing (**c**). (**d**) Number of artery segments per Strahler order. (**e**) Semi-manual segmentation of LCA, highlighting the left coronary circumflex branch (LCx, red). (**f**) Visual representation of branch levels in 3D reconstructed LCx traces. (**g**) Number of branch levels of LCx at each timepoint. Arrowheads indicate rate of change per day. (**h**) Number of arterial (art.) tips in the LCx at each age. Arrowheads indicate rate of change per day. (**i**) Average art. segment length in LCx at each timepoint. (**a**,**d**) n=1 P6, n=1 P60 hearts; (**b**) n=4 P6, n=3 P25 hearts; (**c**) n=5 P6, n= 4 P60 hearts; (**g**,**h**,**i**) n=3 P0, n=7 P6, n=3 P11, n=2 P18, n=2 P25, n=2 P32 hearts. Scale bars: **f**, P0, 100 μm; P6, 200 μm; P18, 400 μm; P32, 500 μm. Error bars are st dev: **, p≤.01.

If arteries do not increase in diameter, additional branches must be added to at least partially offset the increased flow that accompanies heart growth. We next quantified branching during postnatal development. Comparing the Strahler ordering of the two stages revealed that the number of distal vessels (order 9 and 8) were vastly increased (Fig. 7d), aligning with qualitative observations from imaging (see Fig. 1). Since the 3D SimVascular models did not contain arterioles distal to tertiary branches, we further investigated morphometry by manually tracing all α-SMA vessels in a representative branch—the Left Circumflex (LCx)(Fig. 7e, red). Imaris software filament tracing binned each segment of the LCx according to branching levels and quantified the number of arteriole tips (Fig. 7f). The number of branching levels spiked between P0-6 and then hit a plateau until another spike between P25-32 (Fig. 7g). Number of tips increased linearly up to P18 with another spike between P25-32 (Fig. 7h). The P6-18 plateau in number of branch levels compared to the linear increase in number of tips over the same time period indicated that the coronary arteries grow by adding branch segments along the entire length of existing branches. We also observed that the length of each segment was constant among all ages tested (Fig 7i). This results in a coronary tree with many lateral branch segments of a set length.

### Human fetal and adult coronary collateral arteries

A subset of human hearts contains collateral arteries, which are easily observed during an angiogram and are correlated with increased survival in heart disease patients^55, 56^. We sought to identify how our computational modeling studies could help us better understand human collateral function. Thus, we compared the data available from human hearts to our mouse models. We measured vessel diameters for the collaterals observable in angiograms from five patients living with chronic total occlusions (Fig. 8a). This patient population was chosen because their collaterals would be expected to sufficiently support myocardial perfusion downstream of the occlusion without exercise. To compare to mouse data, we also normalized human diameters to the most proximal segment of the LCA. Collateral diameters were on average 15 percent of the LCA (Table 1). These values were in between those observed in the neonate and adult mouse hearts (Table 1). However, a limitation of this comparison is that diameters in angiograms were measured in a 2D projection, which may affect accuracy of absolute values. We also found an average of 2 collaterals per heart (Table 1), but comparisons with mouse data using this parameter are less desirable because angiograms will only highlight a subset of the collaterals that immunostaining would label. These data provide a foundation to determine re-perfusion benefit, but a very precise understanding in humans will need to consider the different pressure distributions resulting from human specific morphology.

**Figure 8:**
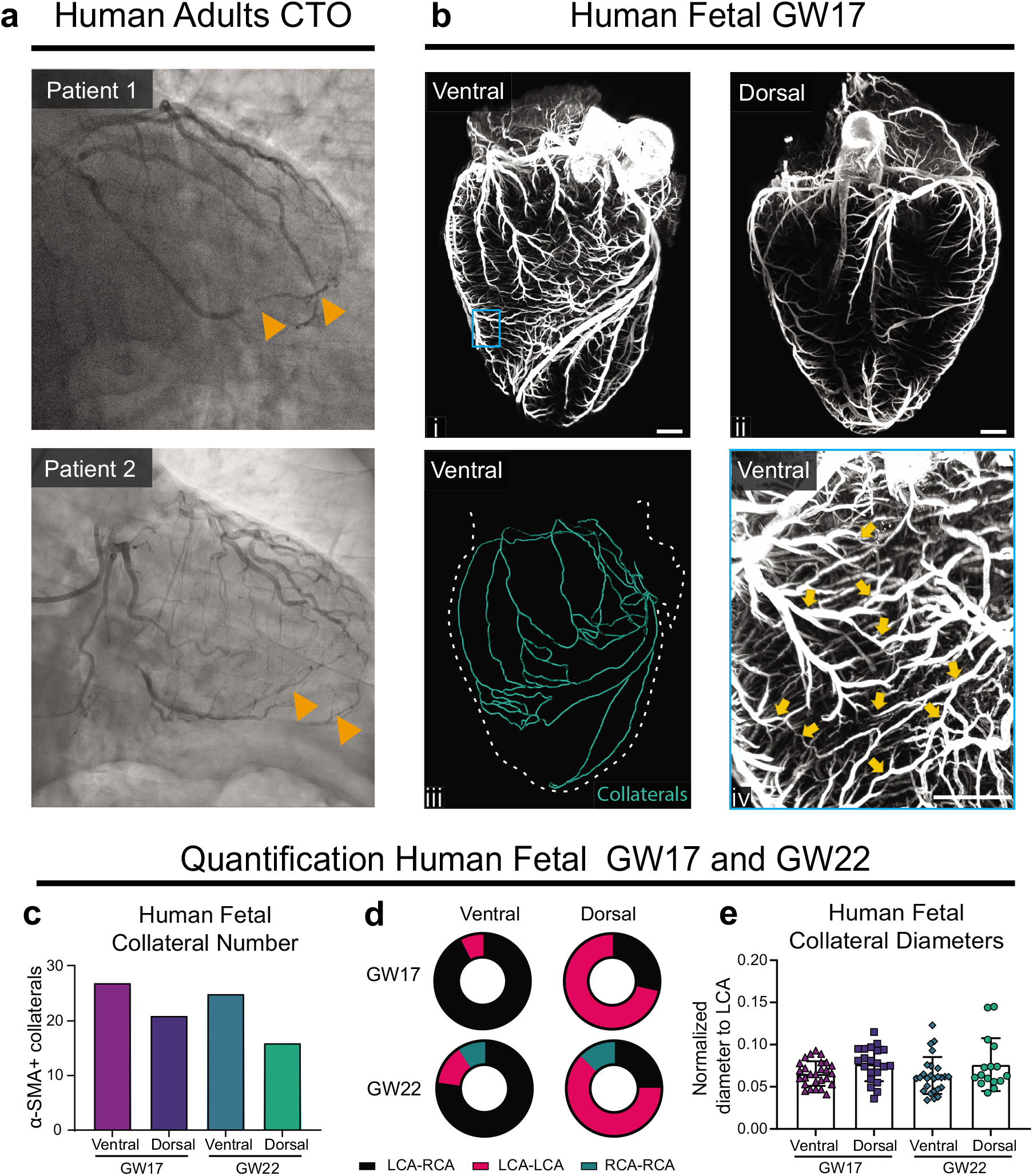
Collateral arteries in adult and fetal human hearts. (**a**) Representative invasive angiograms from adult chronic total occlusion (CTO) patients. Orange arrowheads indicate collaterals. (**b_i-ii_**) Maximum intensity projections of fetal human heart, ventral (**b_i_**) and dorsal (**b_i-ii_**) sides. (**b_iii_**) Traced collateral connections on the ventral side. (**b_iv_**) High magnification (boxed in **b_i_**) of collateral bridges (arrowheads). (**c**-**e**) Quantification of collateral bridge numbers (**c**), connection types (**d**), and diameters (**e**). GW, gestational week. Scale bars: 1 mm.

**Table 1.**
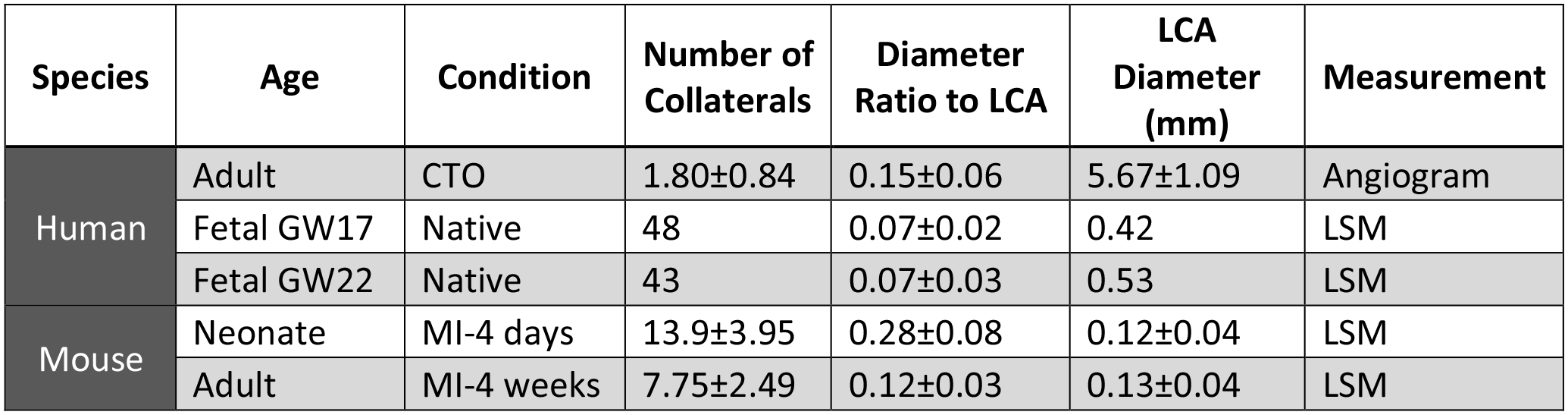
Human fetal hearts have many relatively small collateral arteries. Quantification of collateral artery parameters in human adult CTO patients (n=5), fetal GW17 (n=1) and GW22 (n=1) hearts, neonate (P6) 4 days post-MI (n=9), and adult (16-weeks) 4 weeks post-MI (n=8) mouse hearts. GW, gestation week; P, postnatal day; CTO, chronic total occlusion; MI, myocardial infraction; LCA, left coronary artery; LSM, light sheet microscopy. Reported values are mean ± st dev.

Using post-mortem perfusions, studies from the 1960s reported the presence of coronary collateral arteries in infants and children^57, 58^, but no one has reported whether collaterals develop during embryogenesis. Furthermore, using smooth muscle coverage to identify collateral connections in humans has not been done. We processed two fetal hearts aged 17 and 22 weeks with the same whole-organ immunolabeling method used for murine hearts (Fig. 8b_i__-ii_ and Fig. S5). Both hearts had visible collaterals on the dorsal and ventral sides (Fig. 8b_iii__-iv_ and Fig. S5). Remarkably, >17 collaterals were detected per side (Fig. 8c), which suggests that the whole human heart has at least 40 pre-existing, smooth muscle covered collaterals forming during embryonic development. On the ventral side, most connections bridged distal branches of the RCA and LCA while the majority on the dorsal side connected two LCA branches (Fig. 8d). Collateral diameters were not significantly different across locations or between ages and were on average 7 percent of the most proximal LCA segment (Fig. 8e). Thus, unlike mouse, human hearts have mechanisms in place to form native collateral arteries as part of normal development, which could be the precursors for those that could preserve myocardium downstream of an occlusion.

## Discussion

This study is, to our knowledge, the first-time 3D CFD has been used to quantify hemodynamic forces in the adult and neonatal mouse coronary vasculature. The findings here suggest the possible benefit of promoting the growth of fewer, larger arteries and the natural benefit collaterals have in restoring pressure downstream of a stenosis in neonatal hearts.

Our whole-organ immunolabeling method identifying the structural relationship of collaterals to coronary artery branches suggested that, in mice, the septal artery could play a more critical role in cardiovascular recovery than previously thought. Recent studies utilizing flattened hearts for whole-mount imaging failed to distinguish the SpA from the RCA^9^. Here, tissue clearing and Light sheet microscopy allowed visualization of the intact 3D structure and the complete septal artery, revealing its architectural complexity in healthy hearts. Recent studies have outlined SpA development and proposed that the location of its origin from the aorta could significantly impact cardiac recovery during injury models^59^. Our imaging results suggest a potential route to investigate how positional interactions between these branches could impact vascular repair.

While other studies have performed automatic segmentation and flow modeling of mouse brain and retinal vasculature, there is currently no standard for segmentation and 3D flow simulations in the entire mouse coronary vasculature^35^. We manually segmented over 300 vessels of the adult and over 200 vessels of the neonate coronary network to ensure accurate representation and high model fidelity for fluid simulations using SimVascular. Manual segmentations are currently required because the signal-to-noise ratios, even with high performing antibodies such as anti-α-SMA, are easily recognized by the human eye, but can cause errors in fully automatic segmentations. Methods are in development to improve automation, such as TubeMap, which utilizes machine learning algorithms to produce high fidelity automated segmentations of the brain vasculature^60^. Future work will focus on using or developing similar methods to automate segmentation for cardiac vasculature.

One major advantage of CFD modeling over *ex vivo* measurements of experimental samples is the capability to easily modify one feature, i.e. collateral structure, while keeping all other parameters constant. We tested multiple collateral configurations within the same model to understand the relationship of number, position, and diameter on flow recovery, without potential secondary effects from mouse-to-mouse variations in coronary structure. In this study, we considered values above 30% of non-stenotic perfusion levels as being beneficial. This was based on previous *in vivo* and *in vitro* studies suggesting that myocardial tissue receiving less than 25-30% of baseline flow begins to display measurable signs of cardiac dysfunction. For *in vivo*, when patients were subjected to balloon occlusion of the LCA, only those with greater than approximately 30% coronary flow index maintained normal ST-segments during electrocardiogram^51^. During Langendorff perfusion preparations, heart rate and left ventricle pressures began recovering to normal values above 25% of normal perfusion rates^52^. Simulations demonstrated that increasing diameters or positioning collaterals more proximally allowed them to restore a greater volume of cardiac tissue to this 30% re-perfusion value, more so than increasing numbers of smaller collaterals. We believe that these data are valuable to other scientists in the field studying collateral arteries by giving them more precise guidelines of how tested factors affect collateral flow. With this understanding, physical differences between phenotypes or conditions can be more confidently related to functional differences.

It was surprising that even the most effective collateral configuration modeled in the adult—9col, 40μm—re-perfused only ∼20% of myocardium above the 30% ischemic threshold. This is consistent with studies showing significant scar formation in MI models in mice (permanent coronary ligation), even in the presence of collaterals^10^. This underscores the importance of understanding blood flow through experimentally-induced collateral arteries when considering inducing these vessels as a therapeutic option. Conversely, the virtual collaterals in the neonate with the same characteristics were predicted to have a remarkable ability to shunt flow to the ischemic volume-at-risk—consistent with studies demonstrating the resilience of neonates after total occlusion MI experiments. This, combined with our data that neonates form more numerous and larger collaterals naturally in response to injury, may explain why studies show great recovery post-MI in the neonate via collateral arteries in contrast to adults^9^.

This innate difference in collateral function was attributed to a low pressure drop in the neonatal coronary tree compared to adults due to the increased flow in the adult not being compensated by an equal reduction in vascular resistance. It is important to note that at both ages, the pressure at the tips is approximately 40 mmHg, indicating that our two models were segmented to a similar extent. The gradual, steady decrease in pressure in the adult arises from the more extensive branching observed compared to the neonate, which is in agreement with studies of pig coronary arteries^61^. While we demonstrated there is very little pressure drop in native neonate coronary arteries, the pressure downstream of the 3D model is expected to abruptly drop to capillary levels. The reduction in total coronary resistance from neonate to adult is in concordance with general trends found with µCT measurements of coronary vessels >40 µm in mice aged 1 week to 6 weeks old, but quantitatively much greater in our models^41, 62^. While these studies described the coronary tree morphology with quantitative scaling laws, they were not able to quantify small diameter arteries, which is evident by the noticeably missing arteries in their 1-week mouse coronary model compared to our Light sheet imaged P6 hearts. Future work could investigate how these quantitative scaling laws apply to vessels we are able to visualize with our method.

We next sought to compare our mouse results to human data. It was unexpected that fetal human hearts contained more native collaterals than the injured neonate and adult mouse as well as diseased adult human hearts, although it is difficult to make comparisons between whole-organ α-SMA staining and angiograms. This finding corresponds with data that suggests collaterals tend to decrease during adolescent years because more were found in the fetal stages compared to neonate humans^57, 58^. Since these collaterals are much smaller than those found in adult human diseased hearts via angiogram, it may indicate that small collaterals in the adult heart go undetected. In addition, if we can preserve and enlarge these collaterals in adulthood, they have the potential to greatly improve cardiac perfusion in patients suffering from CAD.

Overall, by combining advanced computational and imaging techniques, a novel connection between collateral flow and native morphological differences, and thus pressure distributions, was established. By bridging these two fields, we uncovered how fundamental coronary morphology changes from embryonic to adult in both mouse and humans affect collateral flow. These findings provide insight into why coronary collateral arteries are better suited for recovering from an injury in young hearts compared to old.

## Limitations

One limitation of this study is that we were not able to measure subject-specific aortic flows and pressures for each mouse coronary vasculature *in vivo* to use for boundary conditions. However, with literature-derived averaged flows scaled to the model size, we expect this to have a very minor effect on the absolute quantities of collateral flow as described here and thus little effect on the overall relative differences between the collateral configurations and ages.

Another limitation is that it is not currently possible to measure the outlet pressure of the coronary tree *in vivo* to validate computational modeling. Due to the small size and inaccessibility of the coronary vasculature in mice, it is challenging to determine *in vivo* flows and pressures at the pre-capillary level. Pressure measurements taken from rabbits and dogs suggest that 50-70% of the pressure is lost when blood reaches the capillary bed^63, 64^. In our study, we tuned the outlet boundary conditions such that the resistance of the 3D model was 65% of the total coronary resistance, which matches expected values from literature^63^. We tested the sensitivity of our results to changes in outlet pressure by adjusting the outlet resistances and found that relative differences between the collateral configurations remained the same.

## Methods

### Animals

All mouse colonies were housed and bred in the animal facility at Stanford University in accordance with institutional animal care and use committee (IACUC) guidance.

### Immunolabeling and iDISCO clearing

Whole heart vasculature staining was performed following the modified iDISCO+ protocol previously described^37, 39^. For all following steps, tissue was always agitated unless noted otherwise. Briefly, animals were perfused with PBS through the dorsal vein, and fixed in 4% paraformaldehyde (Electron Microscopy Science 15714) at 4°C for 1hr (neonatal hearts) or 2hr (adult hearts), washed 3X in PBS and stored in PBS with 0.01% sodium azide (w/v, Sigma-Aldrich S8032) until ready to process. Hearts were dehydrated in increasing series of methanol/ddH_2_O dilutions (20%, 40%, 60%, 80%,100% 2X) for 1hr each, followed by overnight incubation in 66% dichloromethane (DCM, Sigma-Aldrich 34856) and 33% methanol. Next, tissue was washed 2X in methanol 100% for 4hrs and bleached overnight at 4°C in 5% hydrogen peroxide (Sigma-Aldrich 216763) in methanol. Next, the hearts are rehydrated in methanol/ddH-_2_O dilutions (80%, 60%, 40%, 20%) for 1hr each, followed by PBS, 0.2% Triton X-100 PBS (2X) and overnight 20% dimethyl sulfoxide (DMSO), 2.3% Glycine (w/v, Sigma G7126), 0.2% Triton X-100 PBS at 37°C for 2 days. For immunostaining, hearts were blocked in 10% DMSO, 6% Normal Donkey Serum (NDS, Jackson ImmunoResearch 017-000-121) in 0.2% Triton X-100 for 2 days at 37C. Primary antibodies, αSMA-Cy3 conjugated (1:300, Sigma C6198), Connexin-40 (1:300, Alpha diagnostic cx40A), and Podocalyxin (1:1000, R&D Systems MAB1556) were prepared in PBS with 5%DMSO, 32 3% NDS in 0.2% Tween-20, 0.1% Heparin (w/v, Sigma-Aldrich H3393) and incubated at 37°C for 4-14 days. Secondary antibodies conjugated to Alexa 647 (Jackson ImmunoResearch) were matched 1:1 in concentration to their primary target and in prepared in PBS with 3% NDS in 0.2% Tween-20, 0.1% Heparin for the same primary incubation days at 37°C. Washes after each antibody incubation in PBS with 0.2% Tween-20, 0.1% Heparin were performed in 30min increment until the end of the day, followed by an overnight wash. Before clearing, samples were embedded in 1% low-melting agarose (Sigma-Aldrich A9414) in PBS and dehydrated in methanol/ddH_2_O dilutions (20%, 40%, 60%, 80%,100% 2X) for 1hr each and 100% overnight. Next, hearts were incubated in 66% DCM and 33% Methanol for 2.5hrs, followed 2X 30min 100% DCM. Finally, samples were cleared in ethyl cinnamate (ECi, Sigma Aldrich 112372), manually inverted a few times, and kept at RT in the dark until and after imaging.

### Light sheet imaging

Samples were imaged with LaVision BioTec Ultramicroscope II Light sheet microscope in a quartz cuvette filled with ECi. For imaging, we used a MVX10 zoom body (Olympus) with a 2x objective (pixel size of 3.25 µm / x,y) at magnification from 0.63x up to 1.6x. Up to 1400 images were taken for each heart and the z-steps are set to 3.5µm z step size, and light sheet numerical aperture to 0.111 NA. Band-pass emission filters (mean nm / spread) were used, depending on the excited fluorophores: 525/50 for autofluorescence; 595/40 for Cy3; 680/30 for AF647 and 835/70 for AF790. Exposure time was 10ms for single channel and 25ms for multichannel acquisition.

### Perfusion territory mapping

To determine the approximate volume of myocardium each outlet of the coronary model was responsible for perfusing we used (1) a model of the myocardial tissue as the total volume to be perfused and (2) the outlet coordinates as the seed points for the subvolumes. We used the background signal from the staining to segment the model of the myocardial tissue and the cap centers for the outlet coordinates. Then, we used a Voronoi diagram algorithm to assign subvolumes of the myocardial tissue to each outlet of the coronary model such that every point in the myocardial mesh was assigned to the closest outlet. Distances to the closest outlet were determined using a dijkstra algorithm. By integrating the subvolumes of every outlet on each of the 3 main branches (LCA, RCA, and SpA), we were able to calculate the approximate percentage of the total myocardial volume that each main branch is responsible for. We used these percentages as targets for the flow splits when tuning the outflow boundary conditions for the fluid simulations.

We used outlet coordinates instead of centerlines because we were able to better resolve the small coronary arterioles compared to prior studies^65^. This allows us to be certain that myocardial regions close to an outlet are perfused by that outlet, rather than by a large artery nearby that has no outlet nearby.

### CFD simulation

We constructed 3D subject-specific models of the mouse vasculature using SimVascular’s cardiovascular modeling pipeline^44^. Briefly, we created path lines for each vessel (about 349 vessels for the adult and 244 for the neonate). Vessels distal to the quaternary branches were ignored. For each path line, the image data was viewed in planes orthogonal to the tangent of the path line to segment the cross-section. Circles were used to approximate the cross-section, as some areas of the vasculature appeared collapsed or deformed. All segmentations were lofted to create a solid model of each branch, and the branches were then unioned together to form a complete geometric model. Finally, the lofted model was discretized into a linear tetrahedral mesh using the commercial meshing library, MeshSim (Simmetrix, Troy, NY), resulting in a total of 600 thousand and 1.8 million elements for the neonatal and adult models, respectively.

After obtaining the mesh, we uniformly scaled it to account for the shrinkage that occurs via iDISCO. We quantified the volume change due to our specific clearing protocol using water displacement pre- and post-iDISCO and found that the heart shrank to about 63% of its original volume. So, we uniformly scaled the entire volumetric mesh by the inverse (1.58-fold) to ensure that our model faithfully matched the pre-iDISCO geometry.

Inlet boundary conditions were determined as follows. We first determined typical neonate and adult mean pressure and aortic velocity values from literature^45, 46, 66^. Using the mean aortic velocity and the aortic cross sectional inlet area for each mouse used, a subject-specific aortic inflow was calculated and applied. For pulsatile flow simulations, we constructed representative flow waveforms for an adult mouse by digitizing, smoothing, and scaling a waveform from the literature to match the mean inflow at both ages as calculated previously^67^. At the coronary artery outlets, we applied a specialized lumped parameter network to represent the downstream coronary vasculature and the time-varying intramyocardial pressure due to the beating cardiac tissue^48, 49, 68^. The resistance of each coronary outlet was estimated using Murray’s law and tuned such that each of the 3 main branches (LCA, RCA, and SpA) had flow splits equal to the percent volume they perfused. To further tune the capacitances and resistances of the coronary boundary conditions to match literature pressure values^63, 64^, we used a 0D surrogate model for increased efficiency (Fig. S2). At the aortic outlet, we applied a simple RCR boundary condition^47^.

We globally corrected the viscosity in our pulsatile simulations to 1.25cP to account for the Fahraeus-Lindqvist effect; this is necessary because the apparent viscosity of blood decreases in very small tube diameters (<100μm)^69^. While this may significantly underestimate the shear stress in the aorta, the pressure drop in the coronaries was more representative and important for the findings presented here (see Limitations).

We ran blood flow simulations with rigid walls using the stabilized finite element svSolver code in the open-source SimVascular software package^44^ to determine spatially and temporally resolved hemodynamic values, such as pressure, velocity, and wall shear stress at every node in the computational mesh. Simulations ran for 5 cardiac cycles with timesteps of .0001 seconds, and hemodynamic values were determined based on the final cardiac cycle. This took approximately 40 hours on 96 cores via XSEDE and 90 hours on 96 cores via Sherlock. Paraview was used for visualization of the results.

### Virtual collateral placement

Virtual collaterals were strategically added to native coronary vasculature to minimize the initial pressure difference of the two points the collateral was connecting. Specifically, based on an initial simulation of the native vasculature (without any virtual collaterals), a pressure distribution was determined. Using this pressure distribution, virtual collaterals were placed such that each connected equal pressure zones. We replicated realistic connections as closely as possible given size and pressure constraints (Fig. S3). The resistance of each collateral configuration was calculated via Poiseuille’s Law (equation 1).

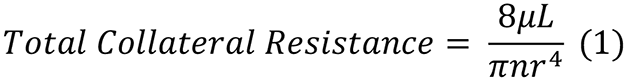

Where µ is the viscosity, *L* is length of the collateral, *n* is the number of collaterals in the configuration, and *r* is the radius of the collateral.

### 3D Resistance

To calculate the resistance of the 3D model, we first generated vessel centerlines via the Vascular Modeling Tool Kit (VMTK; vmtk.org). Each point in the centerline was identified as a branch segment if a perpendicular cross-section at that point did not intersect with any other centerline point. If the cross-section intersected more than one centerline point, then it was labeled as a junction region. This separated the centerline into junctions and branch segments between junctions. After labeling every point, we determined the parent (upstream) branch segment and child (downstream) branch segments for each junction region. We then calculated the resistance for each branch segment based on the pressure difference from the most proximal to distal point and the flow within that segment from the simulation. Finally, the overall 3D resistance was calculated starting from the most distal branches using a recursive method to add the segment resistances in parallel or in series based on the connectivity.

### Diameter-defined Strahler Ordering

We utilized the diameter-defined Strahler ordering system to compare morphometric and hemodynamic quantities at similar positions in the coronary tree between the neonate and adult. This system has been used in previous morphometric studies to classify branch segments into orders that describe the hierarchical nature of a vascular tree^53, 70, 71^. Using the same labels for branch segments and junction regions as in the 3D resistance calculations, we determined the initial Strahler ordering by setting the most distal segments to order 1 and working backwards up the coronary tree to the aorta. Parent segment orders were set to either equal the greater child order if the two children orders were different or incremented by one if the two child orders were the same. Since neither 3D model of the mouse vasculature included all arteries down to the capillary level (only 5 distinct orders here vs. 11 in other studies^54^), we translated all the orders by a constant such that the order of the most proximal segment of the coronaries was 12 and the aorta was order 13 to ensure consistency with previous studies. Segments were then re-organized based on their diameter to ensure that unbalanced branching (i.e. a very small vessel branching from a large one) was properly accounted for. To do this, we iteratively moved segments to higher or lower orders such that every segment within an order was within 1 standard deviation of that order’s mean diameter. With the final diameter-defined Strahler ordering, we compared quantities such as diameter, length, flow, and pressure between the same orders of the neonate and the adult.

### Semi-automated artery tracing

Subsequential images were imported into ImageJ stacks files, these stacks were then converted into 8-bit and resolution reduced to one-fourth the original. Using ImageJ’s plug-in, Simple Neurite Tracer, the branch structures of the LCx were able to be drawn by placing seed points along the length of α-SMA+ vessels^43^. Once every α-SMA+ artery in the LCx branch was completely accounted for within the trace, isolation of the traces was performed by the Fill Out option within the plug-in. The resulting image stack was used as a 3D outline of the arterial structure as the foundation for further modeling and analysis. After discontinuation of Simple Neurite Tracer, the updated version SNT was used in similar manner as above^42^.

### 3D Rendering

The non-traced image stack was overlaid with the filled LCx stack using the Add Channel option in Imaris. Pixel dimensions were updated from the non-reduced 16-bit image metadata. The Filament Object Tracer module was used to generate an Imaris customizable 3D LCx branch model. Branch tips and length were measured by automatically generated data under Number of Terminal Points, and Total Length fields, respectively. Branch levels were obtained from the Filaments Branch Hierarchy field.

Surface objects in Imaris were used for quantifying the sample heart volumes. Myocardium volume was calculated by creating surface objects surrounding the entire sample surface and objects encompassing the lumen of the ventricles. The volumes of the ventricles were then subtracted from the entire heart volume to result in the myocardium tissue volume.

### Murine LCA ligations

Neonatal LCA ligations were performed as previously described^9^ with minimal modifications. P2 neonates were cooled on ice for 6 minutes to induce hypothermic circulatory arrest and placed in a supine position followed by disinfecting with iodine and ethanol. Dissection was carried through the pectoralis major and minor muscles, and the thoracic cavity was entered via the 4th intercostal space. The LCA was identified and ligated at with a doble knot using 8-0 nylon suture, leaving the LCx intact. The chest muscle and skin were then closed (independently) with interrupted 7-0 prolene sutures. The neonate was then allowed to recover at 37°C warm plate and, when conscious, returned to its mother’s care.

Adult mice were performed as previously described^72^. Adult mice were subjected to permanent coronary artery ligation, under anesthesia using initially 1.5%–4% isoflurane chamber for induction. The chest cavity was opened, and a 7-0 silk suture was placed around the left coronary artery, with occlusion verified by blanching of the underlying myocardium. The chest was then sutured closed. Following surgery, Buprenorphine (0.1 mg/kg) was used as an analgesic.

### Immunohistochemistry and confocal microscopy

Neonatal or adult hearts were fixed in 4% PFA overnight at 4 °C, and then cryopreserved in 30% sucrose in PBS for 1 day at 4°C. The following day, coronal heart sections (50 µm in thickness) were cut on a cryostat. Sections were rinsed 3X with PBS, blocked in 5% NDS, 0.5% Triton X-100 in PBS for 1hr at RT, and then incubated with αSMA-Cy3 conjugated (1:300, Sigma C6198) in 0.5% Triton X-100 in PBS overnight at 4°C. Next, sections were rinsed 3X in 0.5% Triton X-100 in PBS and then mounted on slides and covered with Fluoromount G (SouthernBiotech 0100-01). Tissue was imaged using inverted Zeiss LSM-700 confocal microscope at 5x objective. Digital images were captured with Zeiss Zen software and measured using ImageJ.

### Human hearts

Under IRB approved protocols, human fetal hearts were collected for developmental analysis from elective terminations^73^. Gestational age was determined by standard dating criteria by last menstrual period and ultrasound^74^. Tissue was processed within 1hr following procedure. Tissue was extensively rinsed with cold, sterile PBS while placed on ice, followed by incubation in sterile 4% PFA for 4hrs at 4°C before further iDISCO processing. Pregnancies complicated by multiple gestations and known fetal or chromosomal anomalies were excluded.

Human adult samples were acquired from the Stanford Catheterization Angiography Laboratory. All patients displayed symptoms of chronic angina and were scheduled to receive conventional coronary angiography, which was performed according to local clinical standards. Collateral number and size were confirmed by an experienced cardiologist.

### Statistical Analysis

Graphs represent mean values obtained from multiple experiments and error bars represent standard deviation. Unpaired Student’s t test was used to compare groups within an experiment and the level of significance were assigned to statistics in accordance with their p values (0.05 flagged as ∗, 0.01 flagged as ∗∗, less than 0.001 flagged as ∗∗∗, less than 0.0001 flagged as ∗∗∗∗). All graphs were generated using GraphPad Prism software. Error bars represent ± standard deviation.

## Acknowledgements

We thank Andrew Olson and Marco Howard for technical support of Light sheet imaging, and Hanjay Wang for advice on surgical procedures. S.A is supported by BioX Bowes Fellowship. P.E.R.C is supported by the NIGMS of the National Institutes of Health (NIH T32GM007276) and NSF-GRFP (DGE-1656518). M.L.D is supported by the NSF-GRFP (DGE-1656518). D. B. is supported by the Department of Defense CMDRP in Congenital Heart Disease (W81XWH-16-1-0727). K.N. is supported by the NIH/NHLBL (R01HL141712; R01HL146754). A.L.M is supported by NIH (R01EB018302) and NSF Award (1663671). K.R.-H. is supported by the NIH/NHLBL (R01-HL128503) and the New York Stem Cell Foundation (NYSCF-Robertson Investigator).

## Contributions

S.A., P.E.R.C., A.L.M., and K.R.-H. conceived and designed the project. S.A., P.E.R.C., A.N.L.S.-Q., and C.K.C. performed experiments. S.A, P.E.R.C., A.N.L.S.-Q., A.S., and A.M.H. analyzed data. S.A. performed fluid simulations. P.E.R.C., B.C.R., M.Z., and D.B. performed murine cardiac injury studies. K.N. and A.M.P. contributed human adult and fetal samples, respectively. S.A., M.L.D., and M.P. provided analysis tools. S.A. and P.E.R.C. prepared figures. S.A., P.E.R.C., A.L.M., and K.R.-H. wrote the manuscript.

## References

1. Go, A. S. et al. Heart Disease and Stroke Statistics - 2014 Update: A report from the American Heart Association. Circulation vol. 129 (2014).

2. Zimarino, M., D’andreamatteo, M., Waksman, R., Epstein, S. E. & De Caterina, R. The dynamics of the coronary collateral circulation. Nature Reviews Cardiology vol. 11 191–197 (2014).

3. Habib, G. B. et al. Influence of coronary collateral vessels on myocardial infarct size in humans. Results of Phase I thrombolysis in myocardial infarction (TIMI) trial. Circulation 83, 739–746 (1991).

4. Helfant, R. H., Vokonas, P. S. & Gorlin, R. Functional Importance of the Human Coronary Collateral Circulation. N. Engl. J. Med. 284, 1277–1281 (1971).

5. Kim, E. K. et al. A protective role of early collateral blood flow in patients with ST-segment elevation myocardial infarction. Am. Heart J. 171, 56–63 (2016).

6. Meier, P. et al. The impact of the coronary collateral circulation on outcomes in patients with acute coronary syndromes: Results from the ACUITY trial. Heart 100, 647–651 (2014).

7. Red-Horse, K. & Das, S. New Research Is Shining Light on How Collateral Arteries Form in the Heart: a Future Therapeutic Direction? Current Cardiology Reports vol. 23 (2021).

8. Maxwell, M. P., Hearse, D. J. & Yellon, D. M. Species variation in the coronary collateral circulation during regional myocardial ischaemia: A critical determinant of the rate of evolution and extent of myocardial infarction. Cardiovascular Research vol. 21 737–746 (1987).

9. Das, S. et al. A Unique Collateral Artery Development Program Promotes Neonatal Heart Regeneration. Cell 176, 1128–1142.e18 (2019).

10. Zhang, H. & Faber, J. E. De-novo collateral formation following acute myocardial infarction: Dependence on CCR2+ bone marrow cells. J. Mol. Cell. Cardiol. 87, 4– 16 (2015).

11. He, L. et al. Genetic lineage tracing discloses arteriogenesis as the main mechanism for collateral growth in the mouse heart. Cardiovasc. Res. 109, 419– 430 (2016).

12. Porrello, E. R. et al. Regulation of neonatal and adult mammalian heart regeneration by the miR-15 family. Proc. Natl. Acad. Sci. U. S. A. 110, 187–192 (2013).

13. Hara, M. et al. Impact of coronary collaterals on in-hospital and 5-year mortality after ST-elevation myocardial infarction in the contemporary percutaneous coronary intervention era: a prospective observational study. BMJ Open 6, e011105 (2016).

14. Lucitti, J. L. et al. Variants of Rab GTPase-effector binding protein-2 cause variation in the collateral circulation and severity of stroke. Stroke 47, 3022 (2016).

15. Billinger, M. et al. Physiologically assessed coronary collateral flow and adverse cardiac ischemic events: A follow-up study in 403 patients with coronary artery disease. J. Am. Coll. Cardiol. 40, 1545–1550 (2002).

16. Pohl, T. et al. Frequency distribution of collateral flow and factors influencing collateral channel development: Functional collateral channel measurement in 450 patients with coronary artery disease. J. Am. Coll. Cardiol. 38, 1872–1878 (2001).

17. Traupe, T., Gloekler, S., De Marchi, S. F., Werner, G. S. & Seiler, C. Assessment of the human coronary collateral circulation. Circulation vol. 122 1210–1220 (2010).

18. Factor, S. M. et al. Coronary microvascular abnormalities in the hypertensive-diabetic rat. A primary cause of cardiomyopathy? Am. J. Pathol. 116, 9 (1984).

19. Braun, E. J. Intrarenal blood flow distribution in the desert quail following salt loading. Am. J. Physiol. 231, 1111–1118 (1976).

20. He, L. et al. Preexisting endothelial cells mediate cardiac neovascularization after injury. J. Clin. Invest. 127, 2968–2981 (2017).

21. Vasquez, S. X. et al. Optimization of MicroCT Imaging and Blood Vessel Diameter Quantitation of Preclinical Specimen Vasculature with Radiopaque Polymer Injection Medium. PLoS One 6, e19099 (2011).

22. Merz, S. F. et al. Contemporaneous 3D characterization of acute and chronic myocardial I/R injury and response. Nat. Commun. 10, 1–14 (2019).

23. Honeycutt, S. E. & O’Brien, L. L. Injection of Evans blue dye to fluorescently label and image intact vasculature. Biotechniques 70, (2020).

24. Les, A. S. et al. Quantification of hemodynamics in abdominal aortic aneurysms during rest and exercise using magnetic resonance imaging and computational fluid dynamics. Ann. Biomed. Eng. 38, 1288–1313 (2010).

25. Seo, J., Ramachandra, A. B., Boyd, J., Marsden, A. L. & Kahn, A. M. Computational Evaluation of Venous Graft Geometries in Coronary Artery Bypass Surgery. Semin. Thorac. Cardiovasc. Surg. (2021) doi:10.1053/J.SEMTCVS.2021.03.007.

26. Min, J. K. et al. Diagnostic Accuracy of Fractional Flow Reserve From Anatomic CT Angiography. JAMA 308, 1237–1245 (2012).

27. Zhao, S. et al. Patient-specific computational simulation of coronary artery bifurcation stenting. Sci. Reports 2021 111 11, 1–17 (2021).

28. Shad, R. et al. Patient-Specific Computational Fluid Dynamics Reveal Localized Flow Patterns Predictive of Post–Left Ventricular Assist Device Aortic Incompetence. Circ. Hear. Fail. 737–745 (2021) doi:10.1161/CIRCHEARTFAILURE.120.008034.

29. Su, B. et al. Numerical investigation of blood flow in three-dimensional porcine left anterior descending artery with various stenoses. Comput. Biol. Med. 47, 130– 138 (2014).

30. Peiffer, V., Rowland, E. M., Cremers, S. G., Weinberg, P. D. & Sherwin, S. J. Effect of aortic taper on patterns of blood flow and wall shear stress in rabbits: Association with age. Atherosclerosis 223, 114–121 (2012).

31. Lindsey, S. E. et al. Growth and hemodynamics after early embryonic aortic arch occlusion. Biomech. Model. Mechanobiol. 14, 735–751 (2015).

32. Vedula, V. et al. A method to quantify mechanobiologic forces during zebrafish cardiac development using 4-D light sheet imaging and computational modeling. PLOS Comput. Biol. 13, e1005828 (2017).

33. Suo, J. et al. Hemodynamic Shear Stresses in Mouse Aortas Implications for Atherogenesis Materials and Methods Geometry Data Acquisition of the Mouse Aorta. (2007) doi:10.1161/01.ATV.0000253492.45717.46.

34. Shannon, A. T. & Mirbod, P. Three-dimensional flow patterns in the feto-placental vasculature system of the mouse placenta. Microvasc. Res. 111, 88–95 (2017).

35. Bernabeu, M. O. et al. Computer simulations reveal complex distribution of haemodynamic forces in a mouse retina model of angiogenesis. J. R. Soc. Interface 11, (2014).

36. Acuna, A. et al. Computational Fluid Dynamics of Vascular Disease in Animal Models. J. Biomech. Eng. 140, 0808011 (2018).

37. Rios Coronado, P. E. & Red-Horse, K. Enhancing cardiovascular research with whole-organ imaging. Curr. Opin. Hematol. 28, 214–220 (2021).

38. Renier, N. et al. IDISCO: A simple, rapid method to immunolabel large tissue samples for volume imaging. Cell 159, 896–910 (2014).

39. Renier, N. et al. Mapping of Brain Activity by Automated Volume Analysis of Immediate Early Genes. Cell 165, 1789–1802 (2016).

40. Pan, C. et al. Shrinkage-mediated imaging of entire organs and organisms using uDISCO. Nat. Methods 13, 859–867 (2016).

41. Feng, Y. et al. Bifurcation asymmetry of small coronary arteries in juvenile and adult mice. Front. Physiol. 9, 519 (2018).

42. Arshadi, C., Günther, U., Eddison, M., Harrington, K. I. S. & Ferreira, T. A. SNT: a unifying toolbox for quantification of neuronal anatomy. Nat. Methods 18, 374–377 (2021).

43. Longair, M. H., Baker, D. A. & Armstrong, J. D. Simple Neurite Tracer: open source software for reconstruction, visualization and analysis of neuronal processes. Bioinformatics 27, 2453–2454 (2011).

44. Updegrove, A. et al. SimVascular: An Open Source Pipeline for Cardiovascular Simulation. Annals of Biomedical Engineering vol. 45 525–541 (2017).

45. Le, V. P. & Wagenseil, J. E. Echocardiographic Characterization of Postnatal Development in Mice with Reduced Arterial Elasticity. Cardiovasc. Eng. Technol. 3, 424–438 (2012).

46. Huo, Y., Guo, X. & Kassab, G. S. The flow field along the entire length of mouse aorta and primary branches. Ann. Biomed. Eng. 36, 685–699 (2008).

47. Vignon-Clementel, I. E., Figueroa, C. A., Jansen, K. E. & Taylor, C. A. Outflow boundary conditions for 3D simulations of non-periodic blood flow and pressure fields in deformable arteries. Comput. Methods Biomech. Biomed. Engin. 13, 625–640 (2010).

48. Kim, H. J. et al. Patient-Specific Modeling of Blood Flow and Pressure in Human Coronary Arteries. doi:10.1007/s10439-010-0083-6.

49. Tran, J. S., Schiavazzi, D. E., Ramachandra, A. B., Kahn, A. M. & Marsden, A. L. Automated tuning for parameter identification and uncertainty quantification in multi-scale coronary simulations. Comput. Fluids 142, 128–138 (2017).

50. Huang, Y., Guo, X. & Kassab, G. S. Axial nonuniformity of geometric and mechanical properties of mouse aorta is increased during postnatal growth. Am. J. Physiol. - Hear. Circ. Physiol. 290, 657–664 (2006).

51. Seiler, C., Fleisch, M., Garachemani, A. & Meier, B. Coronary collateral quantitation in patients with coronary artery disease using intravascular flow velocity or pressure measurements. J. Am. Coll. Cardiol. 32, 1272–1279 (1998).

52. Stoner, J. D., Angelos, M. G. & Clanton, T. L. Myocardial contractile function during postischemic low-flow reperfusion: critical thresholds of NADH and O _2_ delivery. Am. J. Physiol. Circ. Physiol. 286, H375–H380 (2004).

53. Huang, W., Yen, R. T., McLaurine, M. & Bledsoe, G. Morphometry of the human pulmonary vasculature. https://doi.org/10.1152/jappl.1996.81.5.2123 81, 2123– 2133 (1996).

54. Kassab, G. S., Rider, C. A., Tang, N. J. & Fung, Y. C. B. Morphometry of pig coronary arterial trees. Am. J. Physiol. - Hear. Circ. Physiol. 265, (1993).

55. Wustmann, K., Zbinden, S., Windecker, S., Meier, B. & Seiler, C. Is there functional collateral flow during vascular occlusion in angiographically normal coronary arteries? Circulation 107, 2213–2220 (2003).

56. Meier, P. et al. The collateral circulation of the heart. BMC Med. 2013 111 11, 1–7 (2013).

57. Reiner, L., Molnar, J., Jimenez, F. A. & Freudenthal, R. R. Interarterial coronary anastomoses in neonates. Arch. Pathol. 71, 103–112 (1961).

58. Bloor, C. M., Keefe, J. F. & Browne, M. J. Intercoronary anastomoses in congenital heart disease. Circulation 33, 227–231 (1966).

59. Kolesová, H., Bartoš, M., Hsieh, W. C., Olejníčková, V. & Sedmera, D. Novel approaches to study coronary vasculature development in mice. Dev. Dyn. 247, 1018–1027 (2018).

60. Kirst, C. et al. Mapping the Fine-Scale Organization and Plasticity of the Brain Vasculature. Cell 180, 780–795.e25 (2020).

61. Mittal, N. et al. Analysis of blood flow in the entire coronary arterial tree. Am. J. Physiol. - Hear. Circ. Physiol. 289, 439–446 (2005).

62. Huo, Y. et al. Growth, ageing and scaling laws of coronary arterial trees. J. R. Soc. Interface 12, (2015).

63. Chilian, W. M., Eastham, C. L. & Marcus, M. L. Microvascular distribution of coronary vascular resistance in beating left ventricle. Am. J. Physiol. - Hear. Circ. Physiol. 251, (1986).

64. Nellis, S. H., Liedtke, A. J. & Whitesell, L. Small coronary vessel pressure and diameter in an intact beating rabbit heart using fixed-position and free-motion techniques. Circ. Res. 49, 342–353 (1981).

65. Malkasian, S., Hubbard, L., Dertli, B., Kwon, J. & Molloi, S. Quantification of vessel-specific coronary perfusion territories using minimum-cost path assignment and computed tomography angiography: Validation in a swine model. J. Cardiovasc. Comput. Tomogr. 12, 425–435 (2018).

66. Van Doormaal, M. A. et al. Haemodynamics in the mouse aortic arch computed from MRI-derived velocities at the aortic root. J. R. Soc. Interface 9, 2834–2844 (2012).

67. Hartley, C. J., Reddy, A. K., Michael, L. H., Entman, M. L. & Taffet, G. E. Coronary flow reserve as an index of cardiac function in mice with cardiovascular abnormalities. in 2009 Annual International Conference of the IEEE Engineering in Medicine and Biology Society 1094–1097 (IEEE, 2009). doi:10.1109/IEMBS.2009.5332488.

68. Sankaran, S. et al. Patient-specific multiscale modeling of blood flow for coronary artery bypass graft surgery. Ann. Biomed. Eng. 40, 2228–2242 (2012).

69. Fåhræus, R. & Lindqvist, T. THE VISCOSITY OF THE BLOOD IN NARROW CAPILLARY TUBES. Am. J. Physiol. Content 96, 562–568 (1931).

70. Kassab, G. S., Rider, C. A., Tang, N. J. & Fung, Y. C. B. Morphometry of pig coronary arterial trees. Am. J. Physiol. - Hear. Circ. Physiol. 265, (1993).

71. Dong, M. et al. Image-based scaling laws for somatic growth and pulmonary artery morphometry from infancy to adulthood. Am. J. Physiol. - Hear. Circ. Physiol. 319, H432–H442 (2020).

72. Raftrey, B. et al. Dach1 Extends Artery Networks and Protects Against Cardiac Injury. Circ. Res. (2021) doi:10.1161/circresaha.120.318271.

73. Cunningham, F. G. et al. Abortion. in Williams Obstetrics, 25e (McGraw-Hill Education, 2018).

74. Cunningham, F. G. et al. Prenatal Care. in Williams Obstetrics, 25e (McGraw-Hill Education, 2018).

