## Supplemental Figures for "Blood flow modeling reveals improved collateral artery performance during mammalian heart regeneration"

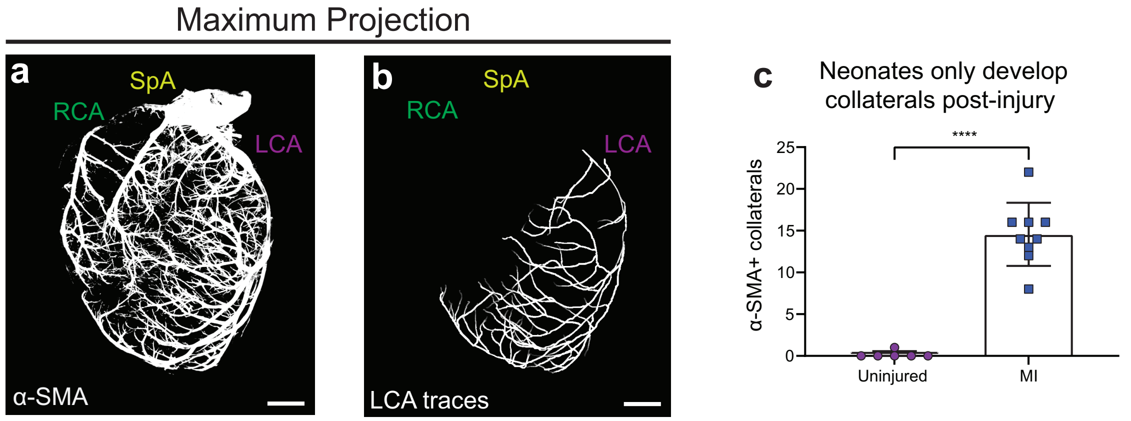


Anbazhakan* and Rios Coronado* et al., Supplementary Figure 1

**Supp Figure 1: Collateral formation in neonatal mice is a response of injury.** (**a**) Maximum intensity projections of representative neonatal P6 intact heart immunolabeled with α-SMA. (**b**) Maximum projection of traces beginning at the most proximal segment of the left coronary artery (LCA) and extended until α-SMA signal discontinue. (**c**) Quantification of collateral numbers in healthy and injured (4-days post MI) neonatal hearts. n=6 uninjured, n=9 injured hearts. Scale bars, 500 µm. Error bars are st dev: ****, p≤.0001.

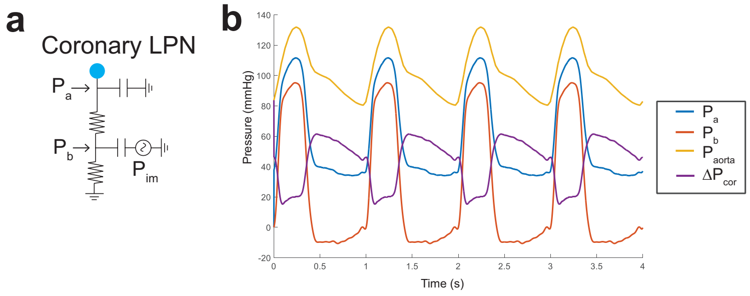


Anbazhakan* and Rios Coronado* et al., Supplementary Figure 2

**Supp Figure 2: 0D surrogate model of pulsatile coronary flow.** (**a**) Schematic of coronary lumped parameter network (LPN). (**b**) Pressure quantities of the coronary LPN. $P_a$, Pressure at point a; $P_b$, pressure at point b; $P_{aorta}$, pressure at aortic inlet; $\Delta P_{cor}$, pressure difference between $P_{aorta}$ and $P_b$.

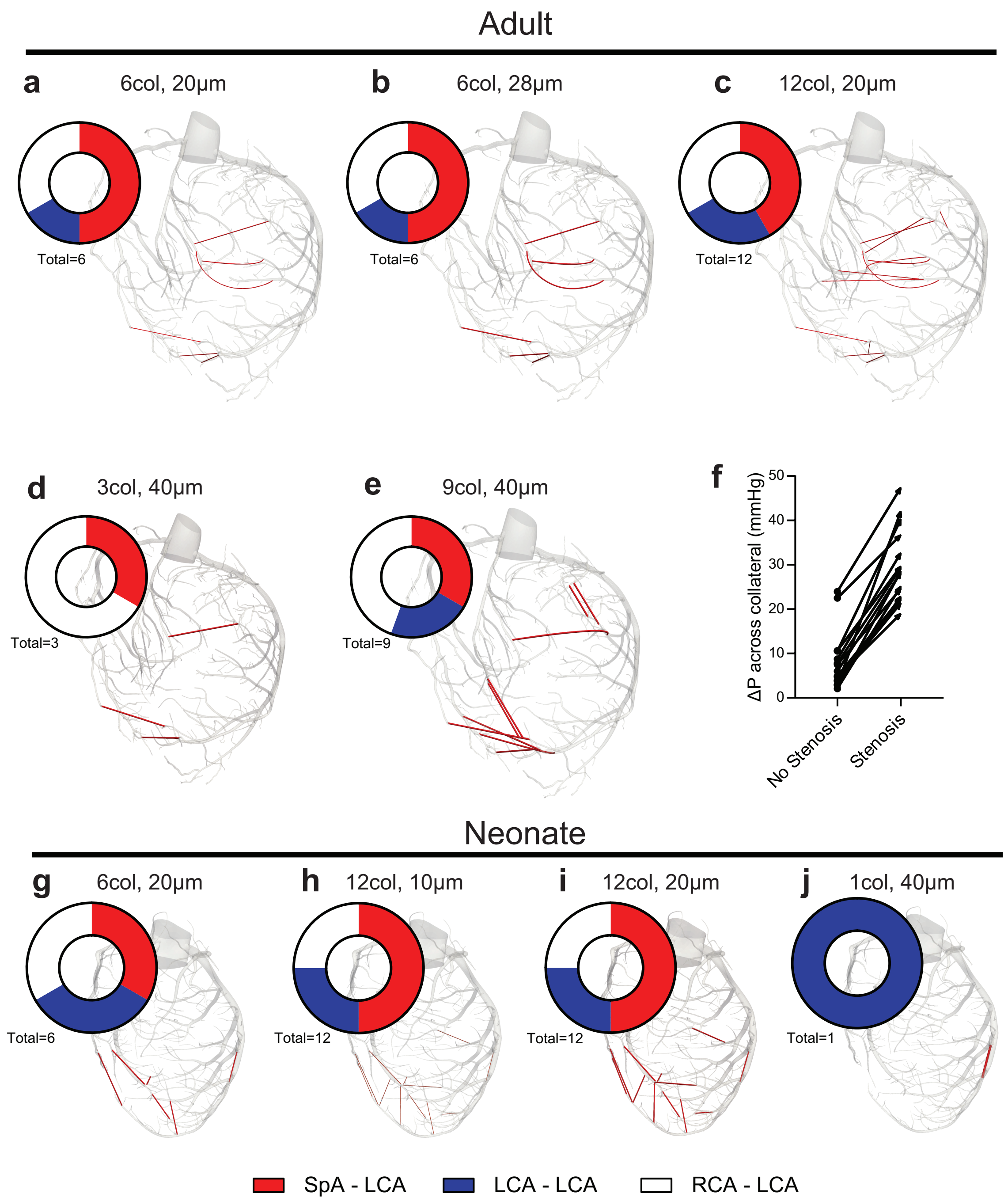


Anbazhakan* and Rios Coronado* et al., Supplementary Figure 3

**Supp Figure 3: Virtual coronary collateral configurations in adult and neonates.** (**a-e**) Adult configurations: 6 collaterals, 20 µm (**a**); 6 collaterals, 28 µm (**b**); 12 collaterals, 20 µm (**c**); 3 collaterals, 40 µm(**d**); 9 collaterals, 40 µm (**e**). (**f**) Pressure increases for each collateral in **c** with and without a 99% stenosis. (**g-j**) Neonatal configurations: 6 collaterals, 20 µm (**g**); 12 collaterals, 10 µm (**h**); 12 collaterals, 20 µm (**i**); 1 collateral, 40 µm (**j**). Pie chart indicates number of collaterals per connection type.

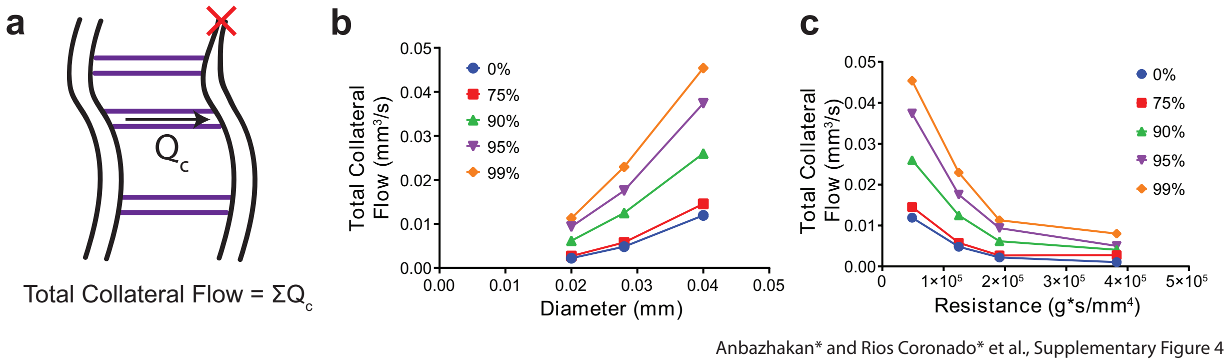


**Supp Figure 4: Verification that collateral flows are in line with expected solutions.** (**a**) Schematic of total collateral flow quantification. (**b**) Total collateral flow vs. diameter of the collateral at each stenosis level. (**c**) Total collateral flow vs. resistance of the collateral configurations 6 collaterals, 20 μm; 12 collaterals, 20 μm; 6 collaterals, 28 μm; 3 collaterals, 40 μm. Resistance was calculated based on the number, diameter, and length of the collaterals via Poiseuille's Law.

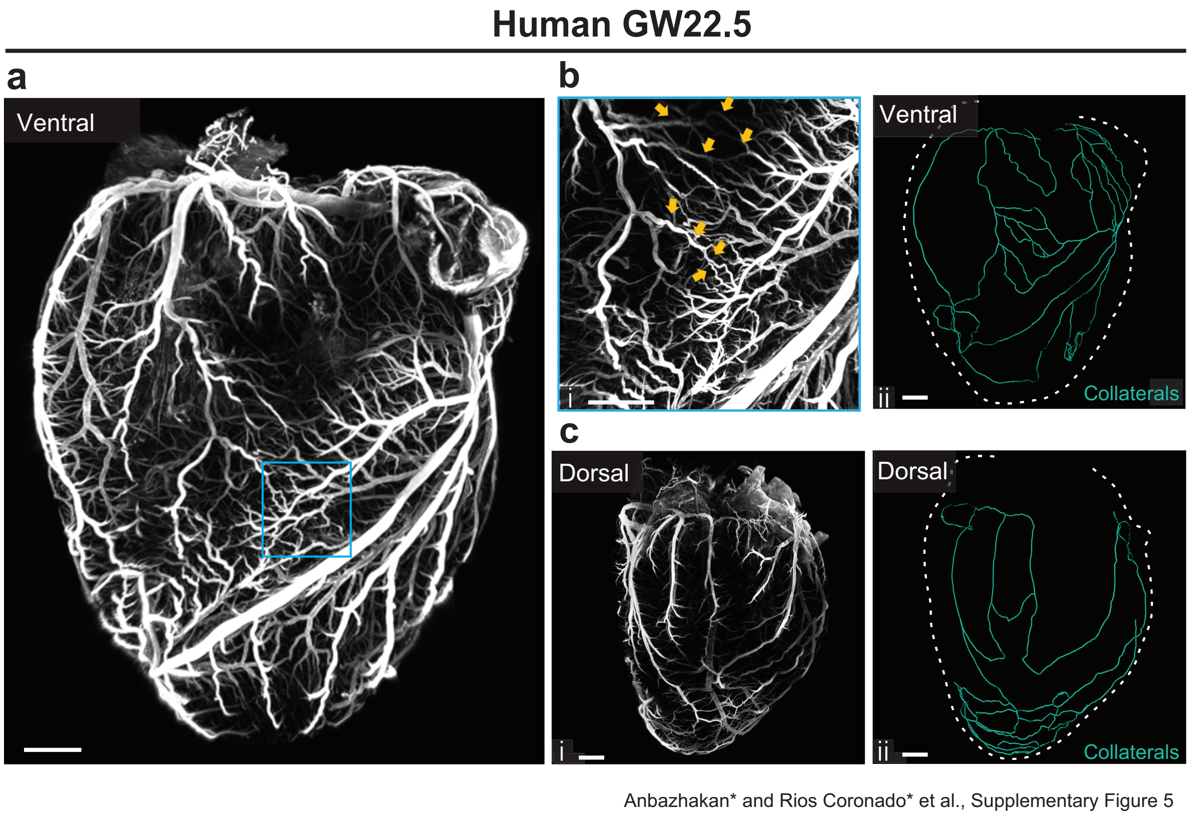


**Supp Figure 5: Collaterals in GW22 fetal human heart.** (**a**) Maximum intensity projection of fetal human heart, ventral side. (**b**$_i$) ROI of ventral side (blue boxed region). Closed orange arrows point to collateral bridges. (**b**$_{ii}$) Traced collateral connection on ventral side. (**c**$_i$) Maximum projection of fetal human heart, dorsal side. (**c**$_{ii}$) Traced collateral connections on dorsal side. Scale bars, **a-c**, 1.5 mm.
